# Mitochondrial competence determines responses to metabolic interventions during aging

**DOI:** 10.64898/2026.05.17.725785

**Authors:** Mingtong Gao, Arshia Naaz, Trishia Yi Ning Cheng, Yizhong Zhang, Sonia Yogasundaram, Nashrul Afiq Faidzinn, Ong Yong Qing Victoria, Raven Rudra Chandramohan, Esther Wong, Rajkumar Dorajoo, Brian K. Kennedy, Mohammad Alfatah

## Abstract

Cellular responses to metabolic interventions vary across physiological contexts, but the basis for this variability remains unclear. Here, we show that mitochondrial competence determines whether cells can engage adaptive metabolic states that support survival during aging. Using 4-methylbenzoic acid (4-MBA), identified as a lifespan-extending compound, as a perturbation probe, we find that modulation of Target of Rapamycin Complex 1 (TORC1) signaling induces a shift from anabolic growth to maintenance-associated metabolism; however, signaling output alone does not predict outcomes. Instead, survival closely correlates with mitochondrial function, with its disruption abolishing adaptive responses. Genetic and biochemical analyses define a mitochondrial regulatory circuit that constrains signaling dynamics and governs state transitions. This regulatory logic is conserved across mammalian systems and operates under oxidative stress, replicative aging, and Hutchinson–Gilford progeria syndrome (HGPS), extending to proliferative cancer cells. These findings establish mitochondrial competence as a key determinant of cellular responsiveness and provide a framework for understanding context-dependent effects of metabolic interventions during aging.

## INTRODUCTION

Cells continuously balance anabolic growth with adaptive programs that sustain survival under conditions of nutrient limitation and stress. This balance becomes particularly critical during aging, where the ability to maintain metabolic homeostasis declines and cellular outcomes increasingly reflect trade-offs between proliferation and long-term persistence. Although pathways that promote growth are well characterized, the mechanisms that govern transitions between growth-associated and survival-supporting metabolic states remain incompletely understood. Defining how such transitions are regulated is essential for understanding variability in responses to metabolic interventions across aging, disease, and stress contexts^1–6^.

The Target of Rapamycin Complex 1 (TORC1) is a central regulator of cellular growth that integrates nutrient availability with environmental cues to drive anabolic metabolism ^7–12^. Reduction of TORC1 activity induces transcriptional and metabolic remodeling programs associated with stress adaptation and maintenance across species ^7,13–15^. However, TORC1 operates within a complex regulatory network shaped by feedback from metabolic flux, organelle function, and stress-responsive pathways, indicating that signaling outputs emerge from network-level integration rather than linear pathway control ^10,16^. How such feedback architectures generate distinct metabolic states that balance growth with survival remains unclear.

Mitochondria are central to this integration. In addition to their roles in energy production and redox homeostasis, mitochondria provide metabolic intermediates and signaling cues that influence cellular state decisions ^17–23^. Bidirectional communication between mitochondria and nutrient-sensing pathways, including TOR signaling, coordinates adaptive responses to environmental and intrinsic stress ^18,22,24–35^. Emerging evidence suggests that mitochondrial state can shape signaling dynamics and influence survival outcomes; however, whether mitochondrial competence acts as a regulatory constraint that determines access to adaptive metabolic programs remains unresolved.

These observations suggest that cells operate within regulated metabolic configurations in which mitochondrial function and nutrient signaling are dynamically coupled to determine whether resources are allocated toward growth or preservation. Conceptual frameworks increasingly propose that cellular behavior reflects transitions between stable regulatory states shaped by network architecture and feedback control ^36^. Defining such configurations requires approaches that capture reciprocal interactions between metabolic state and signaling networks, rather than focusing on individual pathways in isolation.

Building on our previously described TORC1–mitochondria feedback framework (TOMITO) ^37^, we asked whether mitochondrial competence determines the ability of cells to engage adaptive metabolic states during aging. To address this, we leveraged 4-methylbenzoic acid (4-MBA), which we identified as a lifespan-extending compound, as a perturbation probe to interrogate the relationship between nutrient signaling and mitochondrial function. We show that modulation of TORC1 signaling induces a coordinated shift toward a maintenance-associated metabolic state; however, signaling output alone does not predict survival. Instead, mitochondrial competence determines whether cells can execute and sustain adaptive metabolic programs. Through genetic, biochemical, and integrative analyses in yeast, and extension to mammalian systems, we demonstrate that this regulatory logic operates across stress, replicative aging, and Hutchinson–Gilford progeria syndrome (HGPS), as well as in proliferative cancer cells. Together, these findings establish mitochondrial competence as a key determinant of cellular responsiveness and provide a framework for understanding context-dependent effects of metabolic interventions during aging.

## RESULTS

### A small molecule reveals an aging-associated metabolic state that promotes cellular persistence

To examine whether chemical perturbation can uncover mechanisms governing transitions between growth and long-term survival, we investigated the effects of the small molecule 4-methylbenzoic acid (4-MBA) using the yeast chronological lifespan model, a well-established system for studying cellular aging. Across a range of concentrations, 4-MBA exerted modest effects on proliferative growth yet markedly improved maintenance of cell viability over time **(Figures 1A, 1B and S1A–S1F)**. This dissociation between growth and long-term survival indicates that 4-MBA does not act through nonspecific growth inhibition but instead promotes a distinct physiological state associated with enhanced cellular persistence during aging.

**Figure 1.**
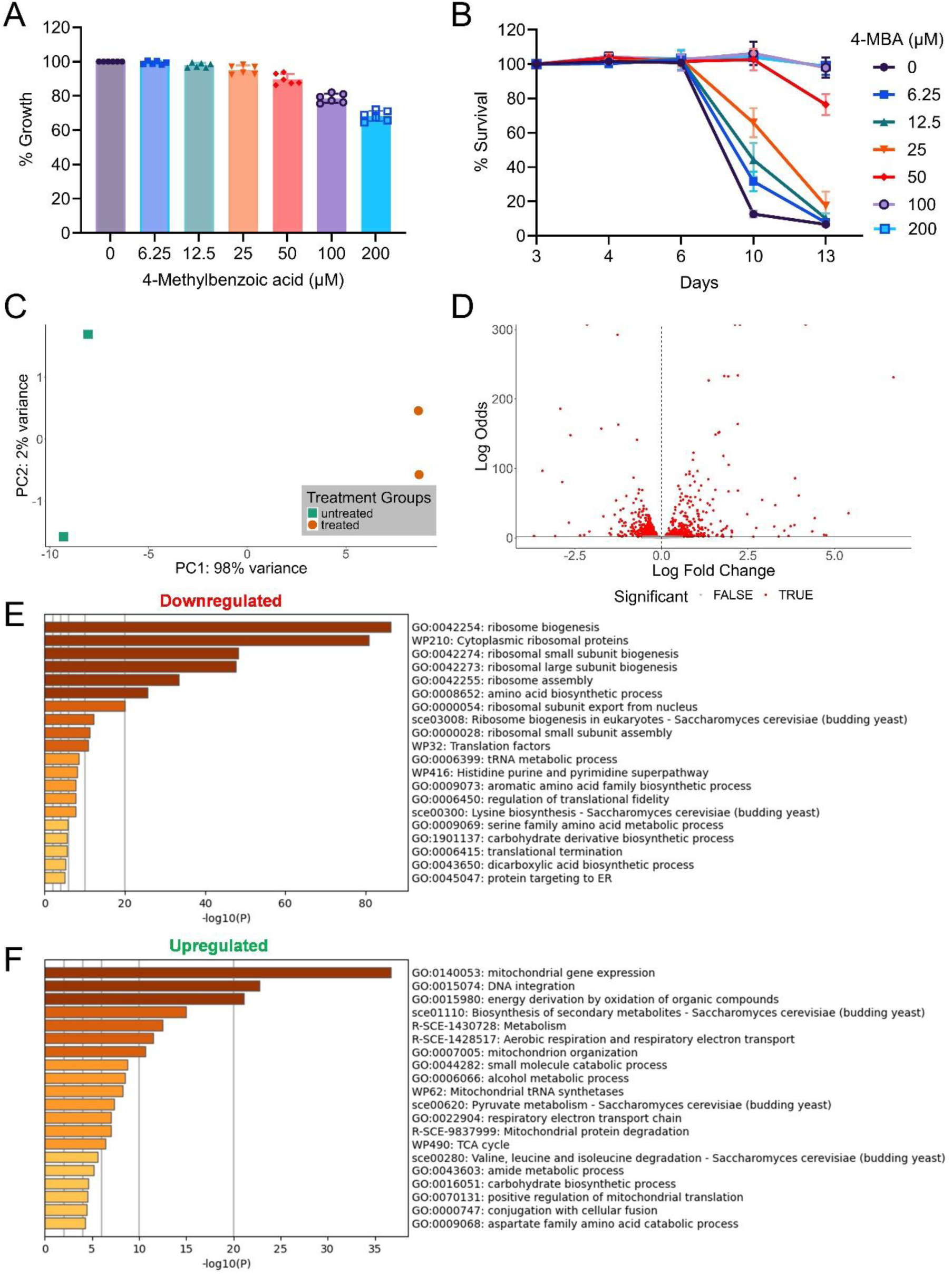
4-Methylbenzoic acid induces an adaptive metabolic state supporting cellular survival (A) Growth of prototrophic *Saccharomyces cerevisiae* CEN.PK113-7D cells cultured in synthetic defined medium in the presence of increasing concentrations of 4-methylbenzoic acid (4-MBA). Growth was assessed at 24 h. Data represent mean ± SD (n = 6). Statistical analyses, including measurements at 48 h and 72 h, are provided in Figure S1A. (B) Chronological survival of cells cultured with increasing concentrations of 4-MBA. Viability at the indicated time points is expressed relative to day 3, which was set to 100% as the first measurement point corresponding to entry into stationary phase. Data represent mean ± SD (n = 6). Statistical analyses for individual time points are shown in Figures S1B–S1E. (C) Principal component analysis of RNA-seq profiles from cells treated with DMSO or 4-MBA (200 μM) for 5 h in synthetic defined medium. Each point represents an independent biological replicate. PC1 and PC2 explain 98% and 2% of total variance, respectively. (D) Volcano plot showing differential gene expression following 4-MBA treatment. Negative and positive log fold changes indicate downregulated and upregulated transcripts, respectively. (E, F) Gene Ontology enrichment analysis of transcripts downregulated (E) or upregulated (F) after 4-MBA treatment. Bar plots show the top enriched biological process categories ranked by −log10(P value).

Maintenance of viability in the absence of proliferation is a defining feature of aging cells. The observed phenotype therefore suggests that 4-MBA induces a metabolic configuration that reallocates cellular resources away from biosynthetic expansion toward maintenance programs that support long-term survival.

To define the molecular basis of this response, we performed global transcriptomic profiling **(Figure 1C and Table S1)**. Principal component analysis revealed clear separation between treated and untreated samples, indicating extensive remodeling of gene expression programs **(Figure 1D)**. Differential expression analysis showed coordinated repression of genes involved in ribosome biogenesis, translation, and amino acid biosynthesis, processes that represent major energetic investments during active growth **(Figures 1E and S1G; Data S1)**. In parallel, genes associated with mitochondrial metabolism, respiratory function, and cellular energy production were induced **(Figures 1F and S1H; Data S2)**.

Together, these changes reflect a redistribution of cellular resources from anabolic growth toward reinforcement of mitochondrial metabolism and maintenance pathways. This coordinated remodeling is consistent with the emergence of an adaptive metabolic state that enhances cellular persistence and decouples proliferative capacity from survival, a hallmark of aging-associated physiological states.

### 4-MBA engages TORC1 through noncanonical regulation

Because repression of biosynthetic programs is a hallmark of reduced TORC1 signaling ^7^, we asked whether the cellular response to 4-MBA reflects modulation of TORC1 activity. Comparative transcriptomic analysis revealed strong concordance between gene expression changes induced by 4-MBA and those triggered by rapamycin, including coordinated downregulation of ribosome biogenesis and other anabolic programs together with induction of metabolic pathways **(Figures 2A, 2B and S2A-S2D; Table S1; Data S3 and S4)**.

**Figure 2.**
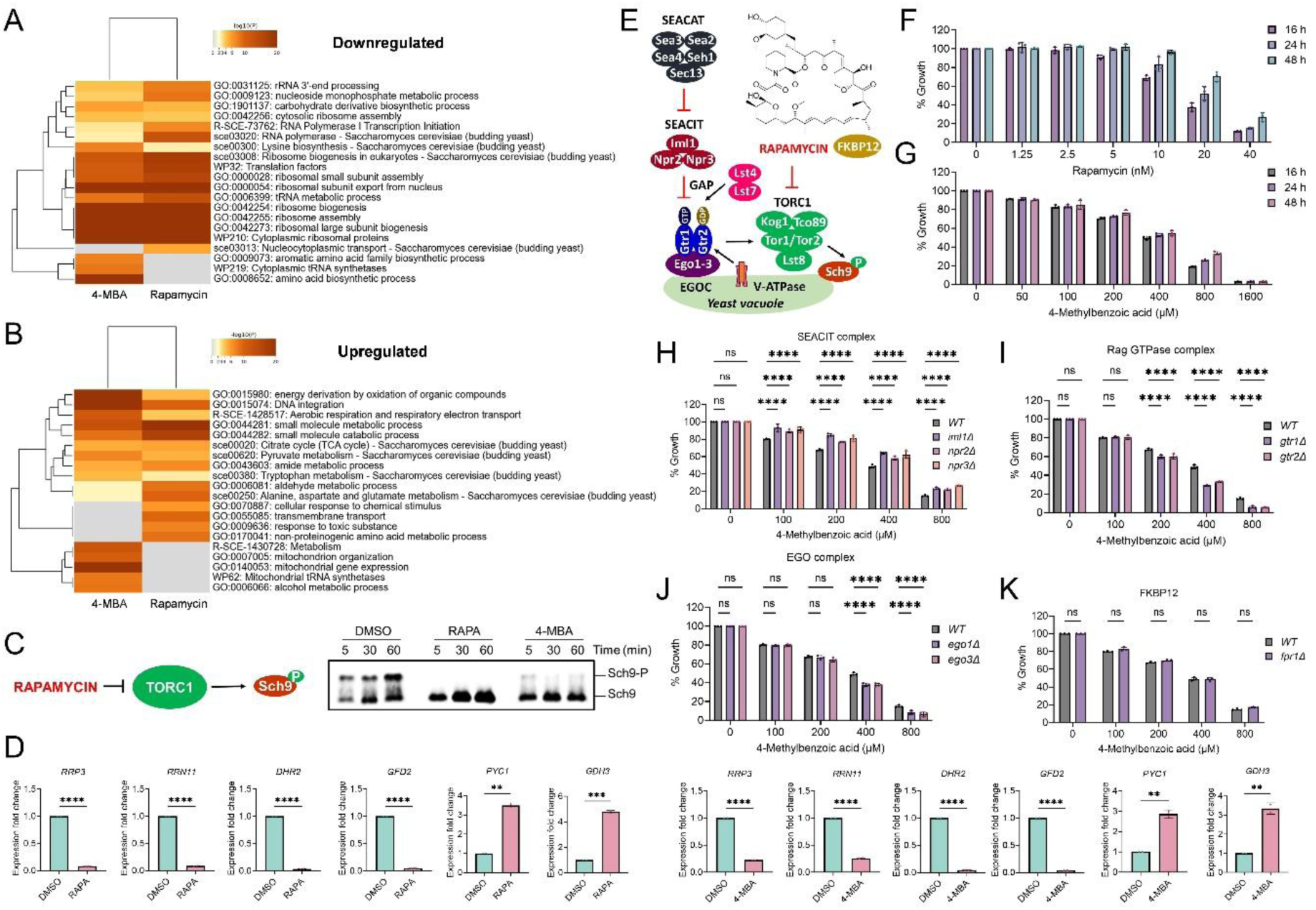
4-Methylbenzoic acid engages TORC1 through a noncanonical mechanism (A, B) Heatmaps comparing functional enrichment profiles of genes downregulated (A) and upregulated (B) following treatment with 4-MBA or rapamycin. Color intensity indicates−log10(P value), and hierarchical clustering highlights similarities and differences in pathway organization between perturbations. (C) Immunoblot analysis of TORC1 activity. Exponentially growing wild-type cells were treated with DMSO, rapamycin (200 nM), or 4-MBA (200 μM), and phosphorylation of the TORC1 substrate Sch9 was assessed at the indicated time points. (D) Quantitative RT–PCR analysis of TORC1-responsive genes following 1 h treatment with rapamycin or 4-MBA. Expression levels were normalized to DMSO-treated controls. Data represent mean ± SD (n = 2). Statistical significance was determined using two-sided Student’s t-test; **P < 0.01, ***P < 0.001, ****P < 0.0001. (E) Schematic representation of TORC1 regulatory pathways in yeast, including the Rag GTPase–EGO axis and SEACIT–SEACAT regulatory modules. (F, G) Growth responses of prototrophic *Saccharomyces cerevisiae* CEN.PK113-7D cells treated with increasing concentrations of rapamycin (F) or 4-MBA (G). Growth is expressed relative to untreated controls. Data represent mean ± SD (n = 3). (H–K) Growth assays of wild-type and TORC1 regulatory mutants treated with increasing concentrations of 4-MBA for 16 h, including SEACIT components (*iml1Δ*, *npr2Δ*, *npr3Δ*) (H), Rag GTPases (*gtr1Δ*, *gtr2Δ*) (I), EGO complex components (*ego1Δ*, *ego3Δ*) (J), and FKBP12 (*fpr1Δ*) (K). Data represent mean ± SD (n = 3). Statistical significance was assessed by two-way ANOVA with Dunnett’s multiple comparisons test; ****P < 0.0001; ns, not significant.

Biochemical analysis showed that 4-MBA reduced TORC1 activity, as evidenced by decreased phosphorylation of the downstream substrate Sch9, comparable to the effect of rapamycin **(Figure 2C)** ^38^. qRT-PCR analysis of canonical TORC1 targets further confirmed a transcriptional response consistent with TORC1 attenuation **(Figure 2D)** ^39^, indicating engagement of TORC1 regulatory outputs.

To position 4-MBA within TORC1 regulatory circuitry, we examined genetic determinants controlling TORC1 activation ^40–46^. Under our experimental conditions, wild-type cells displayed broadly similar growth responses to 4-MBA and rapamycin, supporting convergence on shared growth-regulatory pathways **(Figures 2E-2G)**. Disruption of Rag GTPase or EGO complex components enhanced sensitivity to 4-MBA, whereas deletion of SEACIT components conferred resistance, mirroring established TORC1 regulatory behavior **(Figures 2H-2J and S2E-S2G)**. These interactions indicate that cellular responses to 4-MBA depend on canonical TORC1 regulatory modules. To determine whether 4-MBA acts through the same mechanism as rapamycin, we tested dependence on FKBP12 (Fpr1), which is required for rapamycin-mediated TORC1 inhibition ^47,48^. As expected, deletion of FPR1 abolished responses to rapamycin, yet 4-MBA remained active in *fpr1Δ* cells **(Figures 2K and S2J)**, demonstrating that modulation of TORC1 outputs occurs independently of FKBP12. This regulatory behavior was preserved across genetic backgrounds, with Rag/EGO mutants remaining sensitive and SEACIT mutants resistant **(Figures S2K and S2L)**.

Together, these results show that 4-MBA interfaces with canonical TORC1 regulatory networks while operating through a noncanonical mechanism, establishing TORC1 modulation as a component of the response but not sufficient to fully account for the observed aging-associated phenotype.

### Mitochondrial competence gates engagement of an aging-associated adaptive state

To determine how modulation of TORC1 signaling by 4-MBA reshapes cellular metabolism, we examined transcriptional and functional responses associated with mitochondrial metabolism, a central node linking nutrient signaling to cellular maintenance ^17,28–30,32,33^. Transcriptome analysis revealed coordinated upregulation of genes encoding components of the tricarboxylic acid cycle, electron transport chain, and oxidative phosphorylation, indicating reinforcement of pathways supporting respiratory metabolism and energetic stability **(Figures 1F and S1H; Data S2)**. These changes are consistent with a shift toward a maintenance-oriented metabolic configuration ^15,27^.

To assess whether this remodeling reflects entry into a physiologically relevant adaptive state, we compared transcriptional responses induced by 4-MBA and rapamycin with gene expression profiles from stationary phase cultures, a condition characterized by metabolic adjustment and increased reliance on respiration ^11,27,37,49,50^. Both perturbations showed substantial overlap with stationary phase signatures, including enrichment of mitochondrial pathways and repression of biosynthetic programs **(Figures S3A-S3H; Table S1; Data S5 and S6)**, indicating that chemical modulation recapitulates features of an endogenous adaptive metabolic state.

To independently validate these observations, we quantified expression of representative mitochondrial genes by qRT-PCR. 4-MBA induced transcriptional responses resembling those observed upon TORC1 attenuation, including increased expression of respiratory genes across multiple complexes **(Figures 3A, 3B, S4A and S4B)**. Notably, expression of the oxidative metabolism regulator *HAP4* was elevated, whereas expression of the TORC1-linked mitochondrial regulator *YBR238C* was reduced, indicating reorganization of mitochondrial regulatory networks toward a maintenance-associated state ^37,51,52^. Functionally, cells exposed to 4-MBA displayed increased mitochondrial membrane potential, elevated ATP levels, and increased mitochondrial DNA abundance, indicating enhanced bioenergetic capacity and reinforcement of mitochondrial function **(Figures 3C-3E)**.

**Figure 3.**
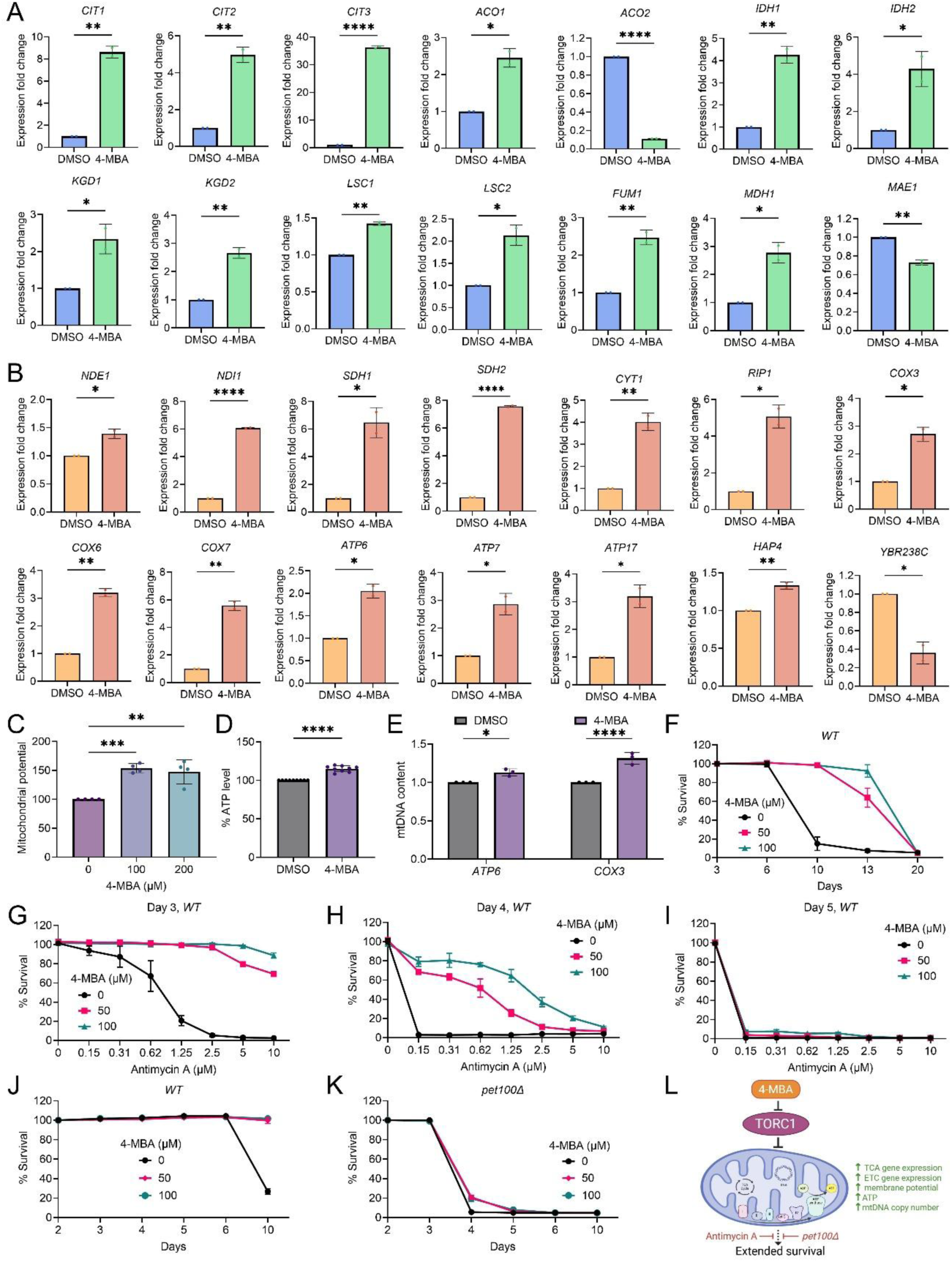
Mitochondrial reinforcement and competence define adaptive metabolic state engagement (A, B) Quantitative RT–PCR analysis of mitochondrial gene expression in prototrophic *Saccharomyces cerevisiae* CEN.PK113-7D wild-type cells treated with 4-methylbenzoic acid (4-MBA; 200 μM) for 1 h. (A) Genes associated with the tricarboxylic acid (TCA) cycle. (B) Genes encoding components of the electron transport chain (ETC) complexes and mitochondrial regulators, including *HAP4* and *YBR238C*. Expression levels were normalized to DMSO-treated controls. Data represent mean ± SD (n = 2). Statistical significance was assessed using two-sided Student’s t-tests; *P < 0.05, **P < 0.01, ***P < 0.001, ****P < 0.0001. (C) Mitochondrial membrane potential in wild-type cells cultured in synthetic defined medium and treated with increasing concentrations of 4-MBA for 24 h. Data represent mean ± SD (n = 4). Statistical significance was determined by one-way ANOVA with Dunnett’s multiple comparisons test; **P < 0.01, ***P < 0.001. (D) Intracellular ATP levels measured in stationary-phase wild-type cells treated with 4-MBA (200 μM). Data represent mean ± SD (n = 9). ****P < 0.0001 by two-sided Student’s t-test. (E) Relative mitochondrial DNA (mtDNA) content in logarithmic-phase wild-type cells treated with 4-MBA (100 μM), quantified by qPCR using mitochondrial genes (*ATP6*, *COX3*) normalized to nuclear *ACT1*. Data represent mean ± SD (n = 3). Statistical significance was assessed by two-way ANOVA with Šídák’s multiple comparisons test; *P < 0.05, **P < 0.01. (F–K) Chronological survival analyses assessing dependence on respiratory function. Wild-type cells were cultured with or without 4-MBA in the presence or absence of the respiratory inhibitor antimycin A or compared with the respiratory-deficient mutant *pet100Δ*. Data represent mean ± SD (n = 3). (L) Proposed model: 4-MBA inhibits TORC1 signaling, promoting mitochondrial remodeling characterized by induction of TCA and ETC gene expression, increased membrane potential, ATP production, and elevated mtDNA copy number. These mitochondrial adaptations are associated with extended survival in a respiration-dependent manner, as antimycin A treatment or disruption of ETC assembly (*pet100Δ*) abolishes the survival benefit.

To determine whether engagement of this state depends on intact respiratory capacity, cells were challenged with the respiratory chain inhibitor antimycin A (AMA) ^53,54^. AMA treatment markedly reduced survival, and neither 4-MBA nor rapamycin restored viability at later timepoints **(Figures 3F–3I and S4C-S4H)**, indicating that the adaptive benefits of TORC1 modulation require functional mitochondria. Consistent with this, 4-MBA failed to support survival in cells with compromised respiratory machinery **(Figures 3J-3L)**.

Together, these transcriptional and functional analyses demonstrate that mitochondrial competence is required for engagement of the adaptive metabolic state that supports cellular persistence during aging. These findings position mitochondrial function as a key determinant of whether TORC1-modulated cells can access survival-supporting states, consistent with a mitochondrial gating mechanism within the TORC1–mitochondria feedback framework (TOMITO) ^37^.

### A mitochondrial regulatory circuit defines the capacity to engage aging-associated adaptive states

To define how mitochondrial regulatory architecture influences adaptive metabolic responses, we examined genetic regulators previously linked to TORC1–mitochondria feedback ^37^. Our previous work identified *YBR238C* as a TORC1-associated factor coordinating mitochondrial function with cellular survival and *RMD9* as a functionally related regulator with opposing effects on viability ^37^. Consistent with these roles, deletion of RMD9 shortened survival whereas RMD9 overexpression extended lifespan, and perturbation of YBR238C similarly altered survival trajectories **(Figure 4A)**, indicating that intrinsic mitochondrial regulatory state influences cellular capacity for long-term persistence during aging.

**Figure 4.**
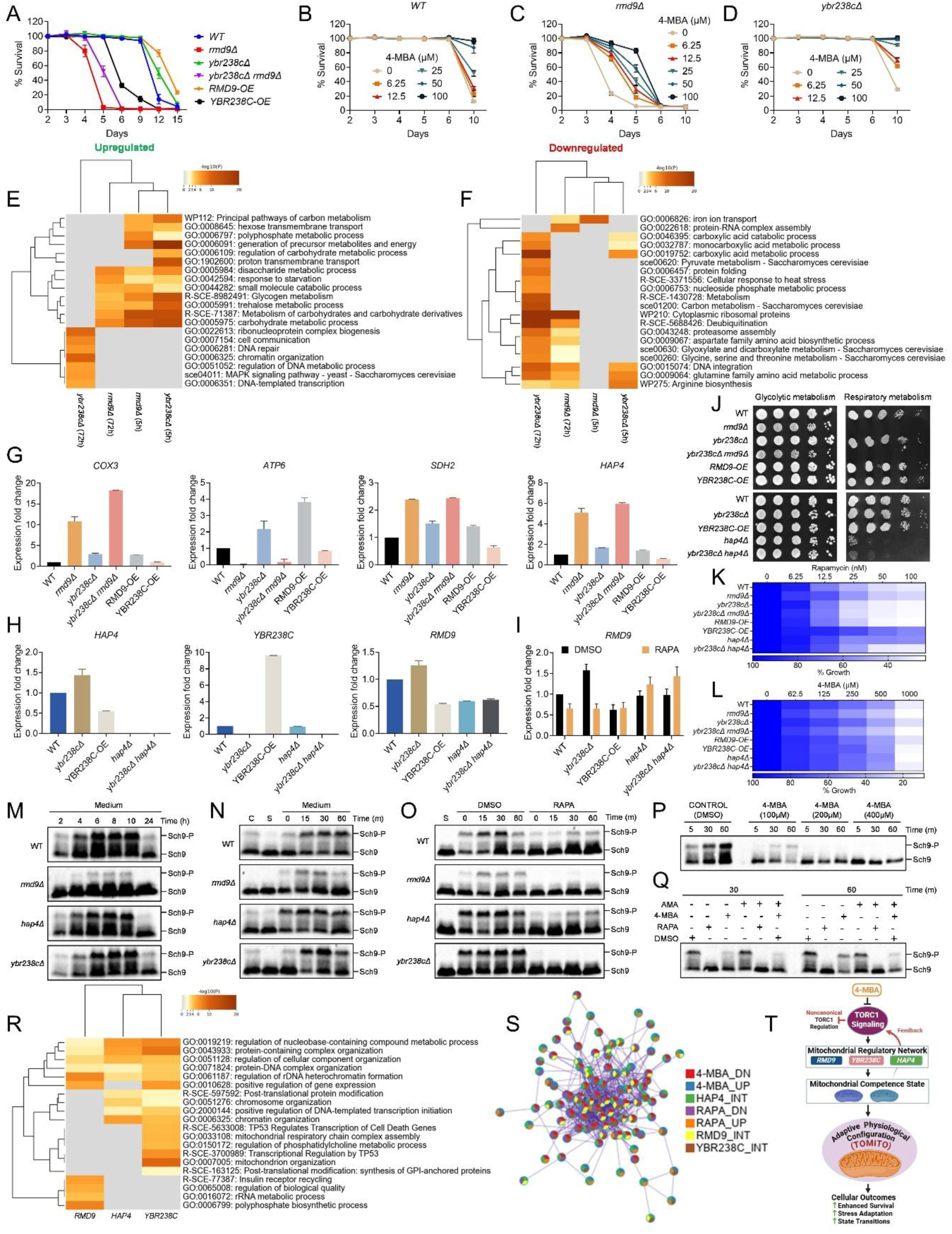
A mitochondrial regulatory circuit defines a competence checkpoint governing signaling dynamics (A) Chronological survival of prototrophic *Saccharomyces cerevisiae* CEN.PK113-7D wild-type cells and mitochondrial regulatory mutants (*rmd9Δ*, *ybr238cΔ*, double mutant, and overexpression strains) cultured in synthetic defined medium. Viability is expressed relative to the initial stationary phase measurement. Data represent mean ± SD (n = 6). (B–D) Dose-dependent survival responses to 4-methylbenzoic acid (4-MBA) in wild-type (B), *rmd9Δ* (C), and *ybr238cΔ* (D) backgrounds. Data represent mean ± SD (n = 3). (E,F) Heatmaps showing functional enrichment of transcripts upregulated (E) or downregulated (F) in *rmd9Δ* and *ybr238cΔ* cells at early (5 h) and late (72 h) time points. Color intensity represents −log10(P value), with hierarchical clustering highlighting similarities and divergence between mitochondrial regulatory states. (G,H) Quantitative RT–PCR analysis of mitochondrial genes (*COX3*, *ATP6*, *SDH2*, *HAP4*) (G) and regulatory factors (*HAP4*, *YBR238C*, *RMD9*) (H) across genotypes during exponential growth. Data represent mean ± SD (n = 2). (I) RMD9 expression following treatment of exponentially growing cultures with DMSO or rapamycin (200 nM) for 1 h across the indicated genotypes. Data represent mean ± SD (n = 2). (J) Serial dilution growth assays on fermentative (glucose) and respiratory (glycerol) media. Representative images from two independent experiments are shown. (K, L) Growth heatmaps summarizing genotype-dependent responses to rapamycin (K) or 4-methylbenzoic acid (4-MBA) (L) across mitochondrial regulatory backgrounds at 48 h. Data represent mean ± SD (n = 2). (M–O) TORC1 signaling dynamics assessed by immunoblotting of Sch9 phosphorylation under basal growth conditions following inoculation into fresh SD medium (M), starvation and nutrient refeeding (C, overnight culture; S, starvation; SD medium refeeding) (N), and rapamycin treatment (O). (P) Dose-dependent effects of 4-MBA on Sch9 phosphorylation over time in exponentially growing cultures. (Q) Effects of mitochondrial respiratory inhibition with antimycin A on TORC1 signaling dynamics in the presence of 4-MBA or rapamycin. (R) Functional enrichment heatmap of physical and genetic interactors of *RMD9*, *YBR238C*, and *HAP4*. (S) Integrated network linking rapamycin-and 4-MBA–induced transcriptomic responses with interaction networks of *RMD9*, *YBR238C*, and *HAP4*. (T) Model depicting a mitochondrial regulatory circuit that establishes a competence checkpoint governing TOMITO engagement and TORC1 signaling dynamics.

Because 4-MBA requires mitochondrial integrity to promote survival, we examined responses across TOMITO genetic contexts. 4-MBA failed to restore survival in *rmd9Δ* cells, indicating that loss of mitochondrial competence limits engagement of the adaptive program, whereas modest enhancement was observed in *ybr238c* mutants **(Figures 4B-4D)**. Transcriptomic analyses revealed that early responses were broadly similar across genotypes but diverged over time, with *YBR238C*-associated states exhibiting progressive transcriptional remodeling while *rmd9Δ* cells remained comparatively static, consistent with impaired adaptive reprogramming **(Figures 4E and 4F; Table S1; Data S7 and S8)**. Notably, transcriptional signatures induced by 4-MBA and rapamycin aligned more closely with *YBR238C*-associated programs, particularly those linked to mitochondrial metabolism **(Figures S5A-S5H; Data S9 and S10)**.

Consistent with these transcriptional patterns, mitochondrial gene expression exhibited genotype-specific remodeling. *RMD9* overexpression drove coordinated induction of respiratory transcripts, whereas *rmd9Δ* displayed a dysregulated profile marked by selective elevation of several mitochondrial genes together with pronounced reduction of *ATP6*, consistent with compensatory activation under conditions of impaired mitochondrial competence **(Figures 4G and S6A)**. Together, these findings indicate that mitochondrial regulatory state determines the capacity of cells to engage adaptive metabolic reprogramming associated with survival during aging.

We next examined regulatory coupling within this network. *YBR238C* perturbation induced *HAP4* expression, and *hap4Δ* exhibited shortened survival **(Figures 4H and S6B)**. *RMD9* expression decreased in *hap4Δ* and *YBR238C* overexpression backgrounds but increased in *ybr238cΔ* cells, and TORC1 inhibition reduced *RMD9* expression in a *HAP4*-dependent manner, revealing reciprocal feedback between mitochondrial transcriptional control and nutrient signaling **(Figures 4H and 4I)**. Functional analyses supported these relationships. *rmd9Δ* and *hap4Δ* mutants displayed respiratory growth defects yet showed resistance to rapamycin, consistent with reduced TORC1 responsiveness under mitochondrial dysfunction, whereas *YBR238C* overexpression conferred resistance to TORC1 inhibition while maintaining respiratory growth **(Figures 4J and 4K)**.

Under 4-MBA treatment, *rmd9Δ* cells remained hypersensitive and deletion of *YBR238C* did not further modify this phenotype, suggesting that *RMD9* defines a key mitochondrial competence checkpoint for chemical adaptation **(Figure 4L)**. Consistent with altered signaling behavior, *rmd9Δ* cells exhibited persistently reduced Sch9 phosphorylation and delayed activation following nutrient refeeding, while *hap4Δ* and *ybr238cΔ* backgrounds displayed distinct phosphorylation dynamics, indicating differential control of TORC1 responsiveness **(Figures 4M-4O)**.

Together, these results define a mitochondrial regulatory circuit in which mitochondrial competence constrains TORC1 signaling behavior and determines the capacity of cells to engage survival-supporting states during aging.

### Mitochondrial feedback shapes TORC1 signaling dynamics and constrains aging-associated outcomes

Given that mitochondrial regulators influence TORC1 behavior and that *rmd9Δ* cells display reduced TORC1 activity together with heightened sensitivity to 4-MBA, we examined how 4-MBA affects TORC1 signaling dynamics. Phosphorylation of the TORC1 effector Sch9 was monitored over time across a range of concentrations ^38^. At higher doses, 4-MBA produced sustained suppression of Sch9 phosphorylation, whereas at lower concentrations it caused a transient decrease followed by recovery, indicating rebound activation of TORC1 signaling **(Figure 4P)**. These dynamics suggest that 4-MBA does not function as a constitutive TORC1 inhibitor but instead engages feedback mechanisms that permit restoration of pathway activity. Consistent with this interpretation, inhibition of mitochondrial respiration with AMA prevented recovery of Sch9 phosphorylation, indicating that mitochondrial integrity is required for dynamic modulation of TORC1 signaling **(Figure 4Q)**.

Comparison with rapamycin revealed distinct signaling behaviors. Rapamycin induced sustained TORC1 inhibition, whereas 4-MBA elicited reversible modulation consistent with feedback regulation rather than direct catalytic blockade, providing a mechanistic basis for the differential responses observed in mitochondrial mutants **(Figures 4M-4O and 4Q)**.

Notably, TORC1 activity alone did not predict survival outcomes; instead, survival closely followed mitochondrial competence across genetic contexts. While 4-MBA transiently reduced TORC1 signaling and enhanced survival, *rmd9Δ* cells maintained low TORC1 activity yet exhibited poor viability **(Figures 4A and 4C)**, with *hap4Δ* showing intermediate phenotypes **(Figure S6B)**. Survival across mitochondrial regulatory backgrounds therefore followed a graded pattern determined by mitochondrial competence.

Network analyses further supported this model. Interactome mapping positioned *HAP4* near *YBR238C* within regulatory networks linking mitochondrial function and nutrient signaling, whereas *RMD9* occupied a distinct module associated with mitochondrial RNA metabolism **(Figure 4R; Table S2; Data S11)**. Integration of these relationships with transcriptomic signatures induced by rapamycin and 4-MBA revealed convergence across regulatory modules, indicating that adaptive responses emerge from coordinated network behavior linking mitochondrial state to TORC1 dynamics rather than TORC1 output alone **(Figure 4S; Data S12)**.

Together, these findings establish that mitochondrial competence governs TORC1 signaling dynamics and constrains the capacity of cells to engage survival-supporting states during aging. Within this framework, mitochondrial state functions as a primary determinant of signaling responsiveness and cellular fate **(Figure 4T)**.

### Adaptive stress responses support aging-associated metabolic states and reveal functional trade-offs

To further define processes that support the adaptive state induced by 4-MBA, we examined stress response pathways associated with mitochondrial function ^27,49,55^. Both rapamycin and 4-MBA induced expression of stress-responsive genes, including targets of the *MSN2/4* transcriptional program and the mitochondrial antioxidant enzyme *SOD2*, consistent with activation of adaptive transcriptional responses accompanying metabolic remodeling **(Figures S6C and S6D)**.

Because mitochondrial reactive oxygen species buffering is essential for maintaining respiratory function, we tested whether antioxidant capacity contributes to cellular persistence. Loss of *SOD2* significantly reduced viability and attenuated the survival benefits conferred by 4-MBA, indicating that mitochondrial antioxidant defenses are required for execution of the adaptive program **(Figures S6E and S6F)**. In contrast, deletion of the mitophagy regulator *ATG32* decreased baseline survival but did not impair responsiveness to 4-MBA, suggesting that mitochondrial turnover is not the primary determinant of engagement of the adaptive state **(Figures S6G and S6H**).

We next asked how this adaptive state influences cellular responses to environmental stress. Cells preconditioned with 4-MBA exhibited increased sensitivity to acute oxidative challenge, indicating that metabolic reprogramming shifts stress tolerance thresholds rather than conferring generalized resistance **(Figures S6I-S6K**).

Together, these findings indicate that the adaptive metabolic state supporting cellular persistence during aging is sustained by mitochondrial antioxidant capacity and is associated with context-dependent trade-offs in stress responsiveness.

### Mitochondria–nutrient signaling coupling underlying adaptive aging-associated states is conserved in mammalian cells

Having established that 4-MBA promotes a coordinated metabolic state transition in yeast, we next asked whether similar regulatory behavior operates in mammalian systems. Because coupling between mitochondrial function and nutrient signaling influences cellular state during aging, conservation would be expected to manifest as coordinated remodeling of signaling and metabolic networks rather than uniform suppression of growth.

We assessed the effects of 4-MBA across multiple mammalian cell types representing distinct metabolic contexts, including C2C12 myoblasts, SH-SY5Y neuronal-like cells, HEK293 embryonic kidney cells, and IMR90 lung fibroblasts. Exposure to 4-MBA produced modest and cell type dependent effects on growth, with greater sensitivity observed in C2C12 and SH-SY5Y cells, whereas HEK293 and IMR90 cells exhibited comparatively mild responses **(Figures 5A–5D)**. Rapamycin suppressed growth across all cell types, consistent with canonical TORC1 inhibition **(Figures S7A–S7D)**. The limited impact of 4-MBA on growth parallels observations in yeast and suggests that its primary effect reflects modulation of cellular regulatory state rather than cytotoxicity.

**Figure 5.**
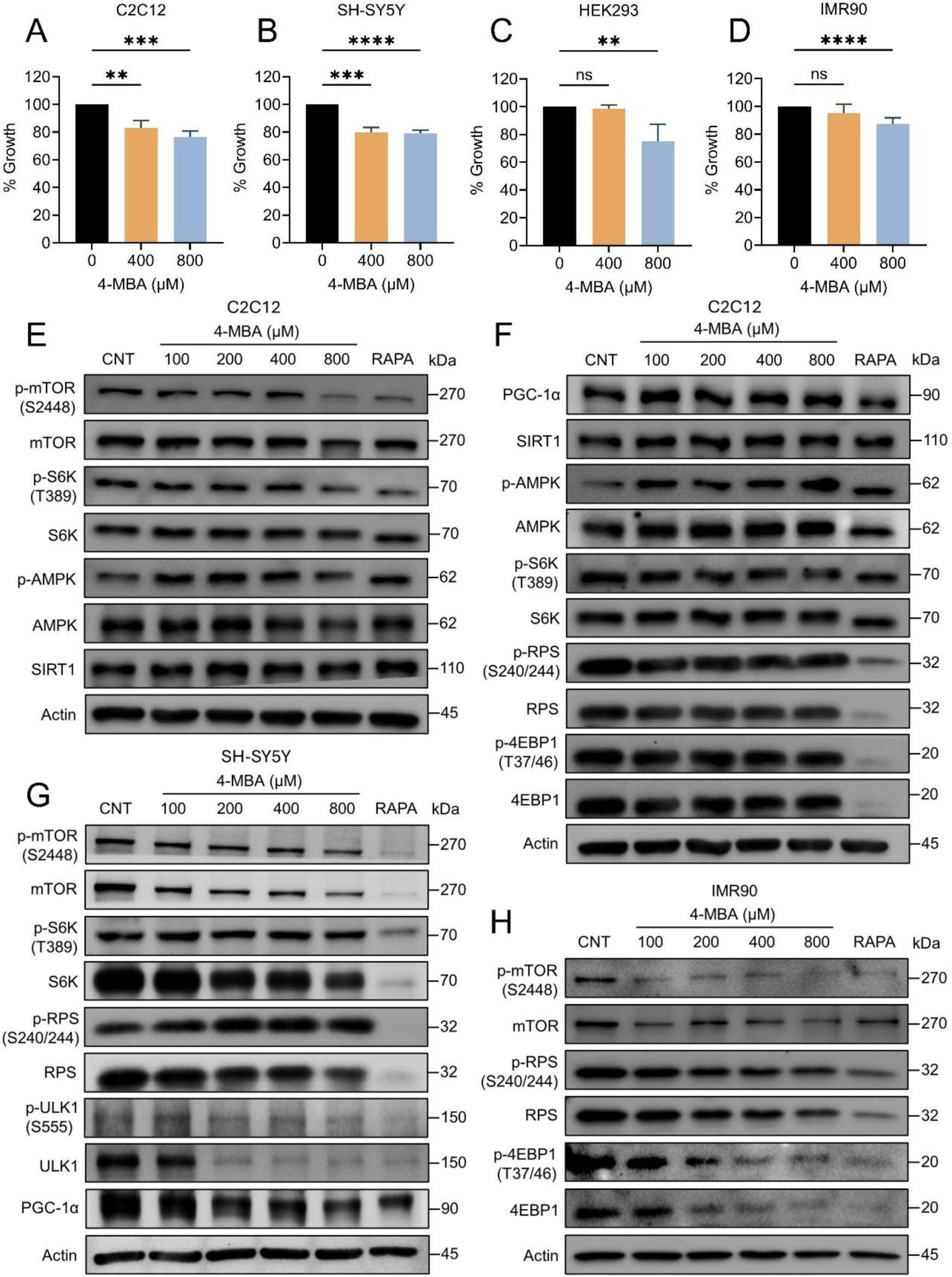
Mitochondria–nutrient signaling coupling is conserved across mammalian cell types (A–D) Growth analysis of mammalian cells treated with 4-methylbenzoic acid (4-MBA) for 48 h, including (A) mouse myoblasts (C2C12), (B) human neuronal-like cells (SH-SY5Y), (C) human embryonic kidney cells (HEK293), and (D) human lung fibroblasts (IMR90). Cell growth was quantified using the CCK-8 assay and is expressed as a percentage relative to DMSO-treated controls. Data represent mean ± SD (n ≥ 3). Statistical significance was determined by one-way ANOVA with Dunnett’s multiple test; **P < 0.01, ***P < 0.001, ****P < 0.0001; ns, not significant. (E–H) Representative immunoblots showing signaling responses in mammalian cells treated with DMSO, 4-MBA (100–800 μM), or rapamycin (100 nM) for 1 h. Cell types include C2C12 (E), SH-SY5Y (F), HEK293 (G), and IMR90 (H). Blots show total and phosphorylated forms of mTOR (Ser2448), S6K (Thr389), ribosomal protein S6 (Ser240/244), and 4EBP1 (Thr37/46), along with phosphorylated and total AMPK, ULK1, SIRT1, and PGC-1α as indicated. β-Actin was used as a loading control.

To determine whether these responses reflect engagement of conserved nutrient-sensing networks, we examined signaling pathways linked to mTOR, AMPK, and mitochondrial regulation ^8,9,17,29,33^. Treatment with 4-MBA produced coordinated modulation of TORC1 signaling, including reduced phosphorylation of canonical downstream targets together with activation of energy-sensing pathways and increased expression of regulators associated with mitochondrial function and metabolic maintenance, including PGC-1α and SIRT1 **(Figures 5E, 5F and S7E)**. These responses closely paralleled those observed with rapamycin, indicating convergence on shared regulatory outputs.

Although the magnitude of signaling changes varied across cell types **(Figures 5G, 5H and S7F–S7H)**, the overall pattern was consistent with engagement of a maintenance-associated physiological configuration characterized by coordinated regulation of nutrient sensing, energy metabolism, and mitochondrial function.

Together, these findings indicate that coordinated coupling between mitochondrial function and nutrient signaling, which supports adaptive metabolic states associated with cellular persistence, is conserved across mammalian systems.

### Reinforcement of mitochondrial capacity underlies adaptive metabolic states in mammalian cells

Because mitochondrial competence is a defining feature of adaptive metabolic states in yeast, we asked whether modulation of nutrient signaling by 4-MBA similarly reinforces mitochondrial capacity in mammalian cells, using rapamycin as a reference for a canonical maintenance-associated state. Treatment with 4-MBA increased mitochondrial DNA abundance, elevated intracellular ATP levels, and enhanced mitochondrial membrane potential across cell types, closely paralleling responses observed following rapamycin exposure **(Figures 6A–6D and S7I–S7K)**.

**Figure 6.**
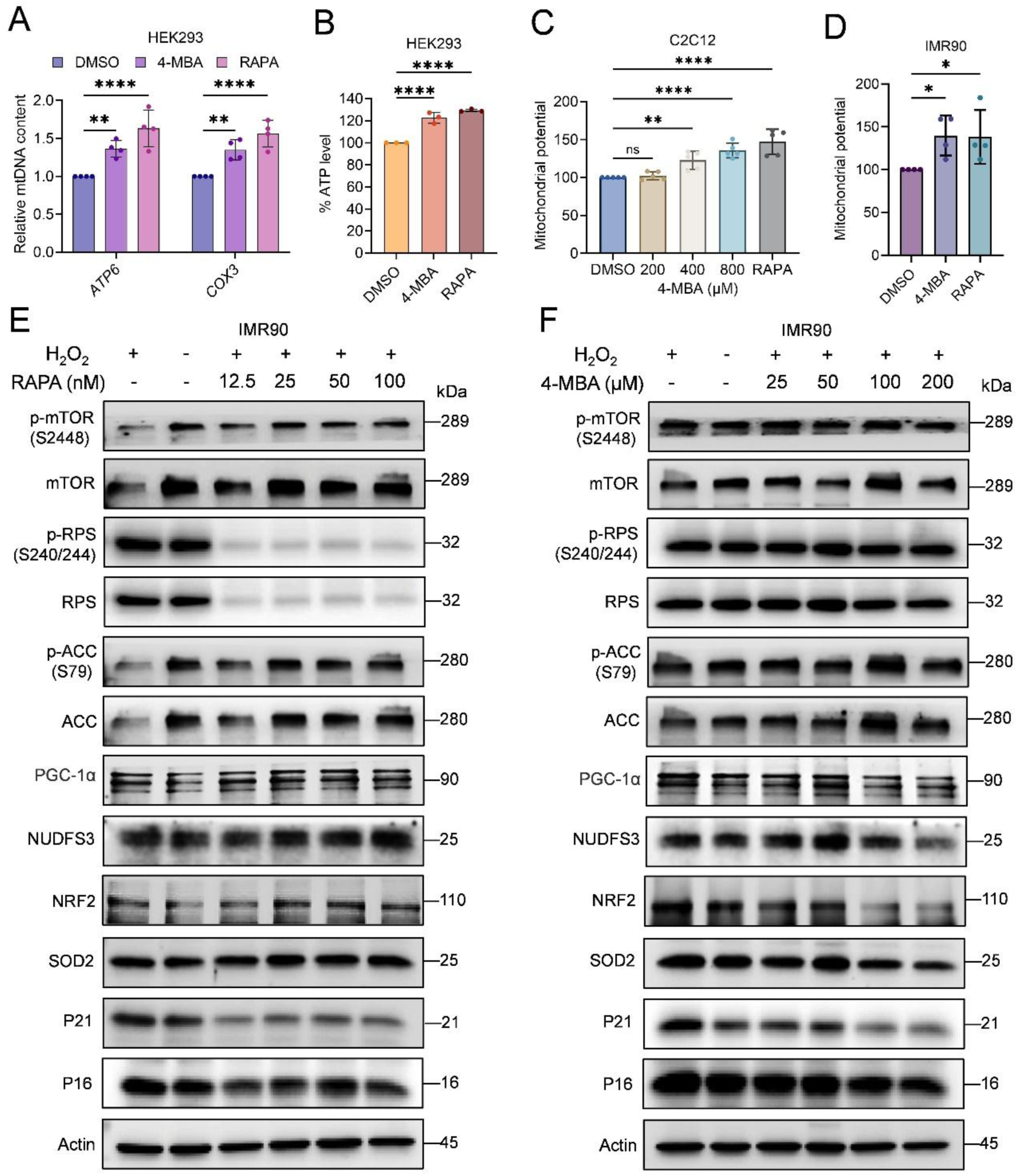
Mitochondrial reinforcement reveals distinct responses to oxidative stress (A) Relative mitochondrial DNA (mtDNA) content in HEK293 cells treated with 4-methylbenzoic acid (4-MBA; 100 μM) or rapamycin (50 nM) for 4 h, normalized to DMSO-treated controls. Data represent mean ± SD (n = 4). Statistical significance was assessed by two-way ANOVA followed by Dunnett’s multiple comparisons test; **P < 0.01, ****P < 0.0001. (B) Intracellular ATP levels in HEK293 cells treated with 4-MBA (200 μM) or rapamycin (50 nM) for 6 h. Data represent mean ± SD (n = 3). Statistical significance was determined by one-way ANOVA followed by Dunnett’s multiple comparisons test; ****P < 0.0001. (C) Mitochondrial membrane potential in mouse myoblasts (C2C12) treated with DMSO, increasing concentrations of 4-MBA (200–800 μM), or rapamycin (100 nM). Data represent mean ± SD (n = 5). Statistical significance was assessed by one-way ANOVA; **P < 0.01, ****P < 0.0001; ns, not significant. (D) Mitochondrial membrane potential in human lung fibroblasts (IMR90) treated with DMSO, 4-MBA (400 μM), or rapamycin (100 nM). Data represent mean ± SD (n = 4). Statistical significance was assessed by one-way ANOVA; *P < 0.05. (E, F) Immunoblot analysis of IMR90 fibroblasts pretreated with rapamycin (E) or 4-MBA (F) for 24 h followed by exposure to hydrogen peroxide (H_2_O_2_; 200 μM) for 4 h. Blots show phosphorylated and total mTOR (Ser2448), ribosomal protein S6 (Ser240/244), acetyl-CoA carboxylase (Ser79), and markers of mitochondrial regulation (PGC-1α, NDUFS3), antioxidant response (NRF2, SOD2), and cell cycle checkpoints (p21, p16). β-Actin was used as a loading control.

These results indicate that 4-MBA promotes reinforcement of mitochondrial bioenergetic capacity, consistent with engagement of a survival-supporting metabolic state associated with cellular persistence during aging. The convergence of mitochondrial responses between 4-MBA and rapamycin indicates that reinforcement of mitochondrial competence represents a conserved feature of adaptive metabolic states across species.

### Oxidative stress reveals distinct modes of adaptive state regulation

To examine how adaptive metabolic states respond to environmental challenge, we exposed IMR90 fibroblasts to hydrogen peroxide to induce a stress-activated state. Oxidative stress triggered coordinated signaling changes, including increased mTOR activity and ribosomal signaling together with induction of mitochondrial regulators such as PGC-1α and NDUFS3 **(Figures 6E and 6F)**. Antioxidant pathways were similarly engaged, as indicated by increased NRF2 and SOD2 levels, accompanied by upregulation of checkpoint markers p21 and p16, reflecting activation of a stress-associated regulatory configuration linking nutrient sensing, mitochondrial adaptation, and cell cycle control.

Treatment with rapamycin suppressed mTOR signaling and reduced checkpoint marker expression while enhancing mitochondrial and antioxidant responses, consistent with a shift toward a maintenance-associated state under reduced anabolic signaling **(Figure 6E)**. In contrast, 4-MBA produced comparatively modest effects on mTOR phosphorylation yet attenuated stress-induced increases in mitochondrial regulators, antioxidant proteins, and checkpoint markers **(Figure 6F)**, indicating selective recalibration of signaling outputs rather than uniform pathway inhibition.

These findings indicate that 4-MBA reshapes stress-induced metabolic states through feedback-dependent modulation of mitochondria–nutrient signaling coupling rather than sustained suppression of TORC1 activity.

Together, these results reveal distinct modes of adaptive state regulation, with rapamycin enforcing sustained suppression of growth signaling and 4-MBA dynamically reprogramming metabolic state through mitochondrial feedback mechanisms.

### Cellular state determines responsiveness of adaptive metabolic programs across stress, aging, and disease contexts

Having observed that oxidative stress induces coordinated remodeling of mitochondria–nutrient signaling networks, we next asked whether similar regulatory configurations arise under intrinsic conditions associated with aging and disease. We therefore examined progeroid fibroblasts and replicative progression as endogenous contexts characterized by chronic stress and altered metabolic regulation ^1,2^.

Progeria fibroblasts exhibited elevated mTOR activity together with increased ribosomal output and induction of mitochondrial regulators including PGC-1α and NDUFS3, accompanied by activation of antioxidant responses marked by NRF2 and SOD2 **(Figures 7A and S8A)**. These changes were associated with increased expression of checkpoint proteins p21 and p16, defining a stress-adapted metabolic state relative to normal fibroblasts, consistent with prior studies linking progeroid syndromes to dysregulated nutrient signaling, mitochondrial remodeling and activation of senescence-associated pathways ^1,2,15,56–60^.

**Figure 7.**
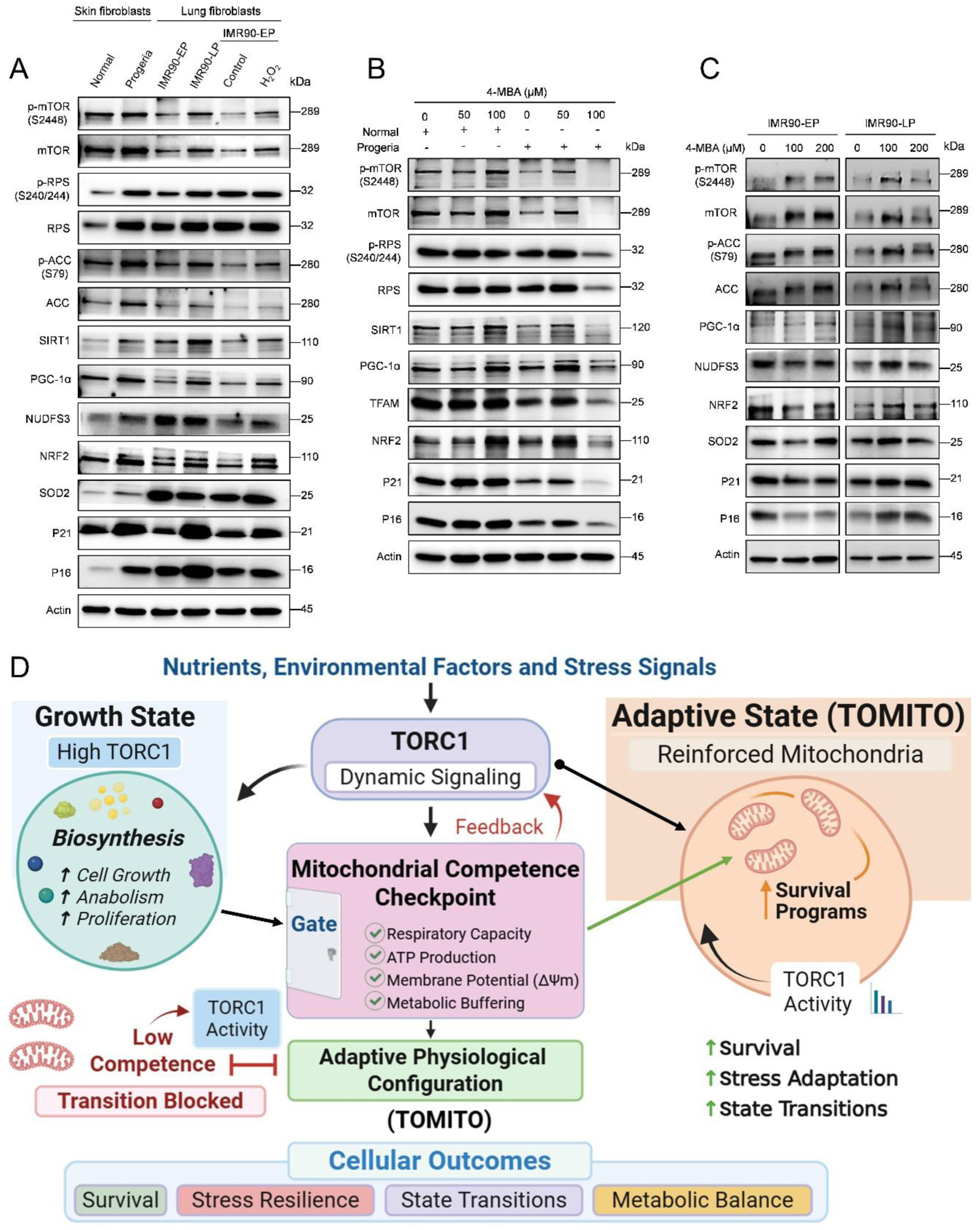
Cellular state determines responses across stress, replicative aging, and progeria (A) Immunoblot analysis comparing signaling states across physiological contexts. Human skin fibroblasts from normal donors (AG03512) and Hutchinson–Gilford progeria syndrome (HGPS; AG03513), both at passage 14, were analyzed alongside human lung fibroblasts (IMR90) at early passage (EP-15) and late passage (LP-25), and early-passage IMR90 cells exposed to hydrogen peroxide (H_2_O_2_; 200 μM, 4 h). Blots show phosphorylated and total mTOR (Ser2448), ribosomal protein S6 (Ser240/244), acetyl-CoA carboxylase (Ser79), and markers of mitochondrial regulation (SIRT1, PGC-1α, NDUFS3), antioxidant response (NRF2, SOD2), and cell cycle checkpoints (p21, p16). β-Actin was used as a loading control. (B) Immunoblot analysis of normal (AG03512; passage 15) and HGPS (AG03513; passage 15) fibroblasts treated with 4-methylbenzoic acid (4-MBA) for 2 h. Blots show phosphorylated and total mTOR (Ser2448), ribosomal protein S6 (Ser240/244), and markers of mitochondrial regulation (SIRT1, PGC-1α, TFAM), antioxidant response (NRF2), and checkpoint proteins (p21, p16). β-Actin served as a loading control. (C) Immunoblot analysis of IMR90 fibroblasts at early passage (EP-16) and late passage (LP-26) cells treated with 4-MBA for 2 h. Blots show phosphorylated and total mTOR (Ser2448), ribosomal protein S6 (Ser240/244), acetyl-CoA carboxylase (Ser79), and markers of mitochondrial regulation (PGC-1α, NDUFS3), antioxidant response (NRF2, SOD2), and checkpoint proteins (p21, p16). β-Actin served as a loading control. (D) Model illustrating mitochondrial competence–dependent control of TORC1 signaling plasticity and adaptive state transitions (TOMITO).

A similar reconfiguration emerged during replicative progression of IMR90 fibroblasts. Late-passage cultures exhibited enhanced mitochondrial and checkpoint signaling compared with early-passage cells, indicating that replicative aging shifts the regulatory balance toward a stress-adapted state **(Figures 7A and S8A)**. Consistent with these observations, early-passage cells exposed to oxidative stress adopted a comparable signaling profile when analyzed alongside progeria and late-passage samples, indicating that acute stress is sufficient to reposition young cells within this adaptive state. Direct comparison across progeria, replicative aging, and acute stress revealed convergence toward a shared regulatory architecture characterized by coordinated nutrient signaling, mitochondrial adaptation, antioxidant activation, and checkpoint engagement.

These findings indicate that diverse aging-associated perturbations converge on a common metabolic state defined by coordinated mitochondrial and nutrient signaling responses.

We next tested whether chemical modulation reshapes signaling behavior across these contexts. Treatment with 4-MBA remodeled pathway outputs in normal and progeria fibroblasts as well as in early-and late-passage IMR90 cells, attenuating stress-associated signaling and recalibrating mitochondrial regulatory programs **(Figures 7B and 7C)**. Importantly, the magnitude and direction of responses varied depending on cellular context, indicating that responsiveness is constrained by underlying physiological state rather than reflecting a uniform inhibitory mechanism.

To determine whether this regulatory logic extends beyond aging-associated fibroblast models, we examined proliferative cancer cells. In MCF7 breast adenocarcinoma cells, 4-MBA similarly induced coordinated remodeling characterized by decreased total mTOR protein abundance, reduced S6K phosphorylation, and induction of mitochondrial regulators PGC-1α and NRF2 **(Figures S9A-S9C)**, indicating that modulation of mitochondria–nutrient signaling coupling reshapes pathway outputs in proliferative contexts.

Together, these findings demonstrate that responsiveness of adaptive metabolic programs is state-dependent across stress, aging, and cancer contexts, with mitochondrial competence acting as a key determinant of signaling behavior and cellular outcomes.

## DISCUSSION

Cellular aging is characterized by a progressive decline in the ability to maintain adaptive metabolic states under conditions of stress and nutrient limitation ^1,2^. Although nutrient-sensing pathways such as TORC1 regulate growth and anabolic metabolism, how cells integrate signaling with mitochondrial function to determine survival outcomes remains incompletely understood ^7–12,16^. Here, we demonstrate that cellular state—defined by mitochondrial competence and metabolic capacity—governs responsiveness to nutrient signaling and determines whether adaptive, survival-supporting metabolic programs can be engaged.

Using 4-methylbenzoic acid (4-MBA) as a perturbation probe, we show that transitions between growth and survival are not dictated by TORC1 inhibition alone but emerge from feedback coupling between nutrient signaling and mitochondrial function. While reduced TORC1 activity has long been associated with lifespan extension through metabolic remodeling programs that enhance respiration and stress adaptation ^27,51,54,55,61^. these models largely position mitochondria as downstream effectors. Our findings extend this framework by demonstrating that mitochondrial competence acts as a regulatory constraint that shapes signaling behavior and determines adaptive outcomes.

A central observation of this study is that modulation of TORC1 outputs induces a coordinated metabolic transition characterized by suppression of anabolic processes and reinforcement of mitochondrial capacity. This shift resembles conserved maintenance-associated programs observed in aging, caloric restriction, and stress adaptation ^7–9,14,15^. However, our data indicate that the mode of TORC1 engagement is critical. Unlike rapamycin, whose inhibitory effects in yeast depend on FKBP12-mediated complex formation and can be bypassed by mutations disrupting this interaction ^47,48^, 4-MBA engages TORC1 regulatory networks independently of FKBP12 and produces reversible signaling dynamics. These findings support the concept that survival-supporting metabolic adaptations can arise through network recalibration rather than sustained suppression of anabolic signaling, consistent with models that describe TORC1 as a metabolic rheostat integrating multiple inputs ^10,16^.

Importantly, our results establish mitochondrial competence as a gating factor controlling access to adaptive metabolic states. Reinforcement of mitochondrial function accompanies engagement of survival-supporting programs, whereas disruption of respiratory capacity abolishes these benefits despite comparable modulation of TORC1 signaling. Thus, signaling output alone does not predict cellular outcomes; instead, survival follows mitochondrial competence. This uncoupling provides a mechanistic explanation for variability in responses to metabolic interventions and suggests that cellular state, rather than pathway activity per se, determines adaptive capacity ^17,30^.

Mechanistically, we identify a mitochondrial regulatory circuit involving *RMD9*, *YBR238C*, and *HAP4* that links organelle function to nutrient signaling behavior. These factors integrate mitochondrial gene expression with nuclear metabolic regulation, establishing a feedback architecture in which mitochondrial status constrains signaling dynamics. This organization is consistent with emerging systems-level models in which cellular behavior reflects transitions between stable regulatory states governed by network architecture rather than linear pathway outputs ^10,16,36^.

The dynamic nature of this regulatory system is highlighted by the observation that 4-MBA induces transient suppression of TORC1 signaling followed by recovery dependent on mitochondrial integrity. In contrast to sustained inhibition by rapamycin, this reversible response suggests that feedback mechanisms restore signaling equilibrium, enabling cells to engage adaptive remodeling without permanently compromising growth capacity. Such dynamics provide a framework for understanding how cells balance growth and maintenance during aging ^47–49,55,61^.

Our findings further demonstrate that adaptive metabolic states involve inherent trade-offs. Engagement of mitochondrial reinforcement and antioxidant programs supports long-term cellular persistence, yet preconditioning alters sensitivity to acute stress, indicating that optimization for survival does not confer uniform resistance. This context-dependent trade-off is consistent with broader principles in aging biology emphasizing the balance between growth, maintenance, and stress responsiveness ^1,2,27,49,55^.

Notably, key features of this regulatory logic are conserved in mammalian systems. Modulation of nutrient signaling by 4-MBA promotes mitochondrial reinforcement and coordinated remodeling of mTOR-associated pathways across multiple cell types. In fibroblasts, similar regulatory configurations emerge under oxidative stress, replicative aging, and progeroid conditions, indicating convergence toward a shared stress-adapted metabolic state. These observations demonstrate that coupling between mitochondrial function and nutrient signaling represents a conserved mechanism governing cellular behavior across aging contexts.

Extending this principle beyond aging-associated models, proliferative cancer cells also exhibit coordinated remodeling of mitochondrial and nutrient signaling pathways in response to 4-MBA. This finding indicates that responsiveness to metabolic interventions is not restricted to senescent or stress-adapted states but is broadly determined by underlying cellular state, reinforcing the generality of this regulatory framework.

Conceptually, our results support a model in which cellular states are defined by network configurations constrained by metabolic capacity rather than by individual signaling pathways. In this framework, mitochondrial competence determines which adaptive states are accessible, thereby governing signaling plasticity and cellular fate. Building on our previously described TOMITO framework ^37^, we show that mitochondrial competence functions as a gating constraint that shapes signaling dynamics and determines adaptive state transitions **(Figure 7D)**. This perspective provides a unifying explanation for context-dependent variability in responses to metabolic interventions, including TORC1-targeting therapies.

Several important questions remain. Although 4-methylbenzoic acid (4-MBA) serves as a useful probe to uncover these dynamics, the extent to which this regulatory logic generalizes across distinct modes and magnitudes of TORC1 modulation remains to be determined. Importantly, the degree and duration of TORC1 inhibition can vary depending on the nature of the perturbation, with lower levels of rapamycin producing partial or transient suppression. Our findings suggest that reduction of anabolic signaling alone is not sufficient to ensure adaptive outcomes; rather, successful engagement of survival-supporting metabolic states depends on mitochondrial competence. Defining how mitochondrial status is sensed and integrated into nutrient signaling networks, and how this coupling interfaces with other hallmarks of aging, will be essential for understanding the broader regulatory landscape governing cellular adaptation.

In summary, we propose that mitochondrial competence acts as a key determinant of cellular responsiveness to metabolic interventions during aging. In this framework, modulation of nutrient signaling creates the opportunity for adaptive metabolic remodeling, but the ability to execute and sustain these transitions is constrained by mitochondrial function. By demonstrating that cellular state governs signaling plasticity through feedback coupling between mitochondria and nutrient-sensing pathways, this work provides a unifying explanation for context-dependent variability in responses to metabolic interventions and establishes a systems-level framework for understanding how growth and survival are coordinated across biological contexts.

## METHODS

### Yeast cultivation, media, and culture conditions

*Saccharomyces cerevisiae* prototrophic CEN.PK113-7D ^62^ and auxotrophic BY4743 (Euroscarf) strains were used in this study. Gene deletion strains were generated using a standard PCR-based gene disruption method ^63^. Yeast strains were revived from frozen glycerol stocks on YPD agar plates (1% Bacto yeast extract, 2% Bacto peptone, 2% glucose, and 2.5% Bacto agar) and incubated for 2–3 days at 30 °C. For experiments, CEN.PK113-7D strains were cultured in synthetic defined (SD) medium containing 6.7 g/L yeast nitrogen base with ammonium sulfate (without amino acids) supplemented with 2% glucose. For auxotrophic BY4743 strains, SD medium was supplemented with histidine (40 mg/L), leucine (160 mg/L), and uracil (40 mg/L).

### Mammalian cell cultivation, media, and culture conditions

C2C12 (mouse muscle–derived), SH-SY5Y (human neuronal-like), HEK293 (human embryonic kidney–derived) and MCF7 (human breast adenocarcinoma) cell lines were obtained from ATCC (USA). IMR90 (human lung fibroblasts), normal human skin fibroblasts (AG03512), and Hutchinson–Gilford progeria syndrome fibroblasts (AG03513) were obtained from the Coriell Institute (Camden, NJ, USA). Cells were cultured in high-glucose Dulbecco’s Modified Eagle Medium (DMEM; Cytiva, USA) supplemented with 10% heat-inactivated fetal bovine serum (FBS; Cytiva, USA) and 1% penicillin–streptomycin (Cytiva, USA). Cells were maintained at 37 °C in a humidified incubator with 5% CO_2_.

### Treatment of cell cultures with chemicals

Cell cultures were treated with rapamycin, 4-methylbenzoic acid, or antimycin A prepared as stock solutions in dimethyl sulfoxide (DMSO). Equivalent volumes of vehicle were added to control samples. The final DMSO concentration did not exceed 1% in yeast experiments and 0.1% in mammalian cell experiments.

### Growth assay

To assess the effects of chemical compounds on yeast growth, assays were performed in a 96-well plate format. *Saccharomyces cerevisiae* prototrophic CEN.PK113-7D strains were cultured in synthetic defined (SD) medium containing 6.7 g/L yeast nitrogen base with ammonium sulfate (without amino acids) supplemented with 2% glucose. For auxotrophic BY4743 strains, SD medium was additionally supplemented with histidine (40 mg/L), leucine (160 mg/L), and uracil (40 mg/L). Yeast cells were inoculated at an initial optical density of approximately 0.2 (OD600) and dispensed in 200 µL per well in 96-well plates containing serial two-fold dilutions of the indicated compounds. Plates were incubated at 30 °C, and growth was monitored by measuring optical density at 600 nm (OD600) using a microplate reader. This assay enabled high-throughput evaluation of growth responses to chemical perturbations

### Chronological aging assay

Cellular aging was evaluated using the chronological lifespan (CLS) assay as described previously ^49,61,64^. *Saccharomyces cerevisiae* prototrophic CEN.PK113-7D strains were cultured in synthetic defined (SD) medium containing 6.7 g/L yeast nitrogen base with ammonium sulfate (without amino acids) supplemented with 2% glucose. To initiate CLS experiments, overnight cultures grown at 30 °C with shaking at 220 rpm were diluted to an initial optical density at 600 nm (OD600) of approximately 0.2 in fresh SD medium. CLS was assessed using two complementary approaches. (i) Microplate outgrowth assay: Cells were aged in 96-well plates containing 200 µL SD medium per well at 30 °C. At indicated time points, 2 µL of stationary-phase culture was transferred into a second 96-well plate containing 200 µL YPD medium and incubated for 24 h at 30 °C. Outgrowth was quantified by measuring OD600 using a microplate reader. (ii) Spot dilution viability assay: Cells were aged in glass flasks containing SD medium at 30 °C with shaking at 220 rpm. At indicated time points, cultures were collected, washed, and normalized to OD600 = 1.0 in YPD medium. Cells were serially diluted 10-fold in YPD, and 3 µL of each dilution was spotted onto YPD agar plates. Plates were incubated for 48 h at 30 °C, and colony outgrowth was documented using a GelDoc imaging system.

### Mammalian cell viability assay

Cell viability was assessed using the Cell Counting Kit-8 (CCK-8; Dojindo, Japan) according to the manufacturer’s instructions. C2C12, SH-SY5Y, HEK293, MCF7 and IMR-90 cells were seeded in 96-well plates and subjected to the indicated treatments for 48 h. Following treatment, 10 µL of CCK-8 reagent was added to each well and incubated at 37 °C. Absorbance was measured at 450 nm using a microplate reader.

### TORC1 activity assay in yeast

TORC1 activity was assessed following established protocols with minor modifications ^65,66^. Prototrophic *Saccharomyces cerevisiae* CEN.PK113-7D wild-type cells expressing Sch9-6×HA were cultured in SD medium and treated under the indicated experimental conditions. Cells were harvested at the specified time points for protein extraction. Proteins were resolved by SDS–PAGE and transferred onto nitrocellulose membranes for immunoblot analysis. Membranes were blocked in 5% (w/v) non-fat milk in TBS containing 0.1% Tween-20 (TBST). Phosphorylated Sch9 was detected using anti-HA 3F10 antibody (1:2000; Roche Life Science), followed by incubation with HRP-conjugated goat anti-rat secondary antibody (1:5000; Santa Cruz Biotechnology). Signals were developed using ECL Prime detection reagent (Amersham) and imaged with an iBright CL1500 imaging system (Thermo Fisher Scientific).

### Western blot

Treated cells were lysed using RIPA buffer (Thermo Fisher Scientific, USA) supplemented with 1x Halt™ Protease and Phosphatase Inhibitor Cocktail (Thermo Fisher Scientific, USA). Protein concentration was quantified using the Pierce™ BCA Protein Assay Kit (Thermo Fisher Scientific, USA), and equal amounts of protein (approximately 20 µg) were prepared with 6x Laemmli buffer (Nacalai Tesque, Japan) containing 5% β-mercaptoethanol (Sigma-Aldrich, USA). Proteins were resolved by SDS-PAGE for 2 hours at 100V, transferred onto nitrocellulose membranes (Bio-Rad, USA) at 85 V for 1.5 hours, and blocked in 1% (w/v) fish skin gelatin (FSG; Sigma-Aldrich, USA) diluted in Tris Buffered Saline-Tween (TBST) for 1 hour. Membranes were incubated overnight at 4°C with primary antibodies against total mTOR (1:1000, CST-2983S) and p-mTOR (Ser-2448) (1:1000,CST-2971), total p70 S6 Kinase (1:1000, CST-9202S) and p-p70 S6 Kinase (Thr389) (1:1000, CST-9234S), total S6 Ribosomal Protein RPS6 (1:1000, CST-2217S), P-S6 Ribosomal Protein RPS6 (Ser-240/244), (1:1000, CST-5364S), total ULK1 (1:1000, CST-8054S) and p-ULK1 (Ser555) (1:1000, CST-5869S), total AMPKα (1:1000,CST-5831S) and p-AMPKα (Thr172) (1:1000,CST-2535S), total ACC (1:1000, CST-3662S) and p-ACC (Ser79) (1:1000, CST-3661S), SIRT1 (1:1000, 13161-1-AP), PGC1α (1:1000, NBP1-04676), TFAM (1:1000, CST-8076S), SOD2 (1:1000, ab13533), NRF2 (1:1000, CST-33649S), p16Ink4a (1:1000, CST-92803S), p21 Waf1/Cip1 (1:1000, CST-2947S), NDUFS3 (1:1000, ab14711), beta-Actin (1:1000, CST; 8457S) followed by incubation with HRP-conjugated secondary antibodies (HRP-linked anti-rabbit IgG (1:3000, CST; 7074S) for 1 hour at room temperature. Detection was performed using Clarity™ Western ECL Blotting Substrate (Bio-Rad, USA) and imaged using the iBrightFL1000 system. Phospho-protein membranes were stripped using Restore™ PLUS Stripping Buffer (Thermo Fisher Scientific, USA).

### RNA extraction and qRT-PCR analysis

Yeast cells were initially mechanically lysed following the manufacturer’s disruption protocol. Total RNA extraction from yeast and human cells was carried out using the Qiagen RNeasy mini kit. The concentration and integrity of RNA were assessed using the ND-1000 UV-visible light spectrophotometer (Nanodrop Technologies). qRT-PCR (Quantitative Reverse Transcription-Polymerase Chain Reaction) experiments were conducted as previously described, utilizing the QuantiTect Reverse Transcription Kit (Qiagen) and SYBR Fast Universal qPCR Kit (Kapa Biosystems) ^65,67^. The abundance of each gene was determined relative to the housekeeping transcript *ACT1* for yeast and *GAPDH* for human cells.

### RNA sequencing

For RNA sequencing, RNA concentration and integrity were assessed using the Bioanalyzer 2100 with the Agilent RNA 6000 Nano Lab Chip kit. High-quality RNA samples were then prepared for paired-end RNA sequencing at the Novogene facility. An enrichment process targeting the polyadenylated mRNA fraction was employed. Following enrichment, complementary DNA (cDNA) libraries were generated from the enriched mRNA via reverse transcription. The quality and size distribution of the cDNA libraries were validated using the Bioanalyzer. RNA sequencing was performed on the NovaSeq PE150 platform. The raw sequencing reads were aligned using the nf-core RNAseq pipeline (nf-core/rna-seq version 3.8.1), which utilized the STAR aligner and RSEM for quantification ^68^. The alignment was done using the *S. cerevisiae* reference genome with corresponding annotations (GTF format, version 1.105) sourced from Ensembl, generating gene-level expression counts ^69^. RNA-sequencing datasets for rapamycin-treated cells, stationary-phase cultures and *ybr238cΔ* cells (5h) were obtained from our previously published reports ^37,50^.

### Functional enrichment analysis

Functional annotation and enrichment analyses were performed using the Metascape online platform ^70^. Physical and genetic interactors of yeast *RMD9*, *YBR238C*, and *HAP4* were retrieved from the Saccharomyces Genome Database (SGD). Enrichment analyses of interactomes and chemical perturbation transcriptomic datasets were conducted using Metascape ^70^, which integrates multiple bioinformatics resources for pathway and process enrichment. Default parameters were applied, and significantly enriched Gene Ontology (GO) terms and pathways were visualized to identify biological processes associated with the datasets.

### Mitochondrial DNA copy number determination

Mitochondrial DNA (mtDNA) copy number was quantified by real-time quantitative PCR (qPCR) as described previously ^67^. Genomic DNA was extracted using the Quick-DNA Midiprep Plus Kit (Zymo Research) according to the manufacturer’s instructions. qPCR reactions (20 μL) contained 20 ng total DNA and SYBR Fast Universal qPCR Master Mix (Kapa Biosystems) and were performed on a QuantStudio 6 Flex system (Applied Biosystems). The thermal cycling program consisted of an initial denaturation at 95 °C for 3 min followed by 40 cycles of 95 °C for 1 s and 60 °C for 20 s. Melt-curve analysis was performed to confirm amplification specificity. Relative mtDNA content was determined by quantifying mitochondrial genes (*ATP6* and *COX3*) and normalizing to nuclear reference genes (*ACT1* for yeast and *GAPDH* for mammalian cells).

### Mitochondrial membrane potential measurement

Mitochondrial membrane potential in yeast was measured using the fluorescent dye DiOC6 as described previously ^71^. Yeast cells were harvested, washed with phosphate-buffered saline (PBS), and stained with 50 nM DiOC6 at 30 °C for 30 min in the dark. Cells were washed to remove excess dye and resuspended in PBS for fluorescence measurement (excitation 482 nm, emission 504 nm). Fluorescence intensity was normalized to OD600. For mammalian cells, cultures were incubated with 50 nM DiOC6 in PBS for 30 min at 37 °C. Fluorescence was measured and normalized to cell viability determined from parallel CCK-8 assays performed on identically seeded and treated plates.

### ATP measurements

ATP levels were measured as described previously ^65–67^. Yeast cells were precipitated with 5% trichloroacetic acid (TCA) on ice for at least 5 min, washed, resuspended in 10% TCA, and lysed using glass beads in a bead beater. ATP was extracted from mammalian cells using Triton X-100 lysis buffer. ATP concentrations were quantified using the PhosphoWorks™ Luminometric ATP Assay Kit (AAT Bioquest) and normalized to total protein content measured using the Bio-Rad protein assay.

## Statistical analysis

All statistical analyses and graphical presentations were performed using GraphPad Prism (v10.6.1). Data are presented as mean ± standard deviation (SD) unless otherwise indicated. Statistical significance was assessed using Student’s t-test, one-way ANOVA, or two-way ANOVA with appropriate multiple comparison corrections as specified in figure legends. Significance thresholds were defined as *P < 0.05, **P < 0.01, ***P < 0.001, and ****P < 0.0001; “ns” indicates not significant.

## Code availability

No custom code or mathematical algorithm was used in this study.

## Data availability

Further information and requests for resources and reagents should be directed to and will be fulfilled by the Lead Contact, Dr. Mohammad Alfatah.

## ACKNOWLEDGMENTS

We thank Sebastian Maurer-Stroh (Executive Director, BII, ASTAR) and Su Xinyi (Acting Executive Director, IMCB, ASTAR) for institutional support and facilitation of research resources. We also thank Prakash Arumugam (SIFBI, A*STAR) for sharing plasmids and Mr. Koh Ting Wei Kelvin (HLTRP, NUS Medicine, Singapore) for assistance with laboratory management. This work is supported by the Young Investigator Research Grant (YIRG), National Medical Research Council, Singapore (MOH-001348-00) and US NAM Healthy Longevity Catalyst Awards Grant (MOH-001439).

## AUTHOR CONTRIBUTIONS STATEMENT

M.G: Investigation, Formal analysis

A.N: Investigation, Formal analysis

T.C.Y.N: Investigation, Formal analysis

Y.Z: Investigation, Formal analysis

S.Y: Investigation, Formal analysis

N.A.F: Investigation, Formal analysis

O.Y.Q.V: Investigation, Formal analysis

R.R.C: Investigation, Formal analysis

E.W: Reviewing and editing

R.D: Reviewing and editing

B.K.K: Reviewing and editing

M.A: Conceptualization, Supervision, Writing-original draft, Writing-review and editing, Funding acquisition.

All authors read, critically reviewed and approved the final manuscript.

M.A is the guarantor of this work.

