## Supplemental Figures for "Mitochondrial competence determines responses to metabolic interventions during aging"

#Equal authorship contributions

*To whom the correspondence should be addressed.
 (Mohammad Alfatah)

**Keywords**
Aging; Mitochondrial competence; TORC1 signaling; Metabolic interventions; Cellular state transitions; Stress adaptation; Hutchinson–Gilford progeria syndrome (HGPS); Yeast–mammalian conservation

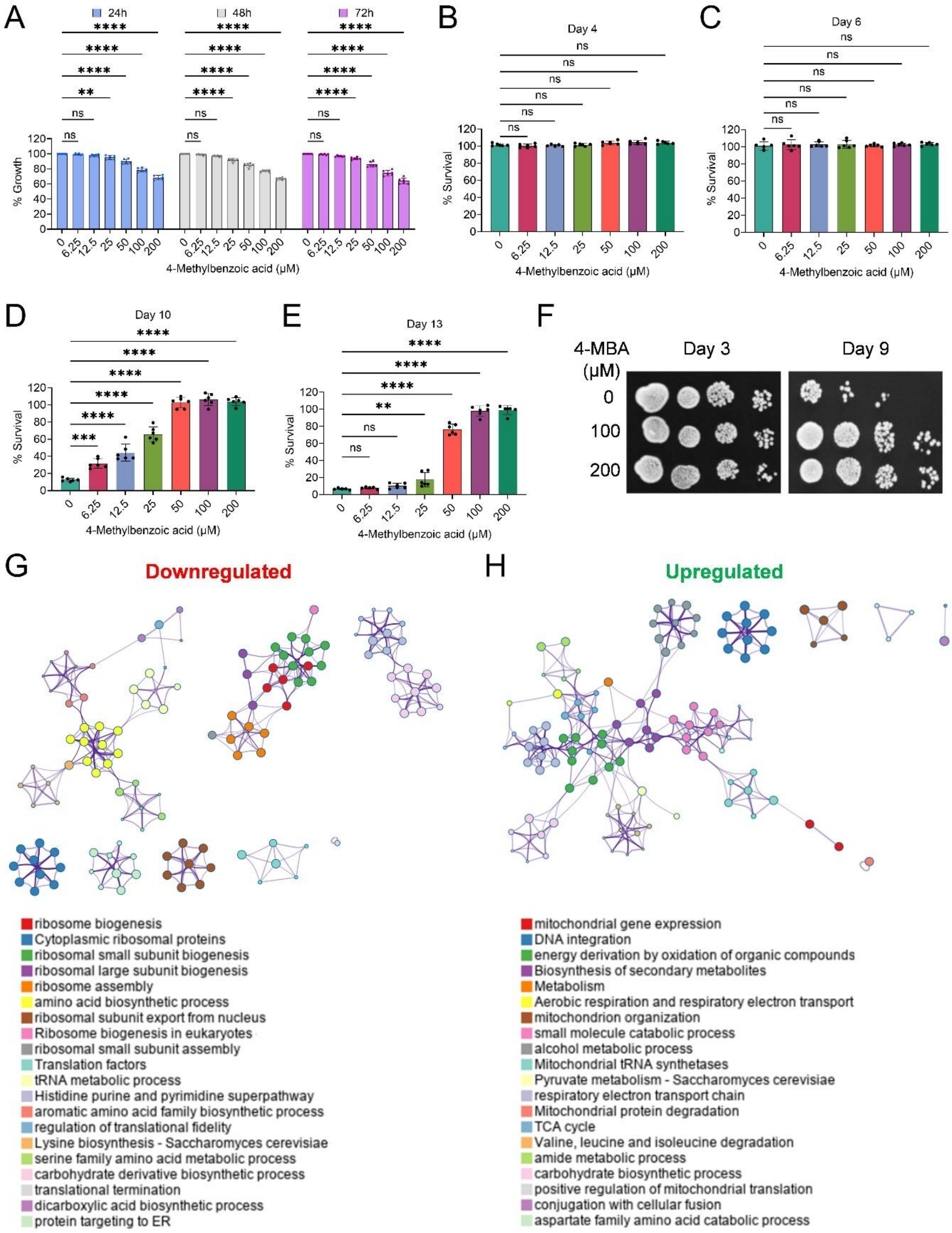


**Figure S1. 4-Methylbenzoic promotes survival with minimal effects on growth and induces transcriptional remodeling, related to Figure 1**

(A) Growth of prototrophic *Saccharomyces cerevisiae* CEN.PK113-7D cells cultured in synthetic defined medium supplemented with increasing concentrations of 4-methylbenzoic acid (4-MBA). Growth was measured at 24 h, 48 h, and 72 h. Data represent mean ± SD (n =

6). Statistical significance was assessed by two-way ANOVA with Dunnett's multiple comparisons test; $**P < 0.01$, $****P < 0.0001$; ns, not significant.

(B–E) Chronological survival of cells treated with increasing concentrations of 4-MBA measured at day 4 (B), day 6 (C), day 10 (D), and day 13 (E). Viability at each time point is expressed relative to day 3, which was set to 100% as the first measurement following entry into stationary phase. Data represent mean ± SD (n = 6). Statistical significance was assessed by one-way ANOVA with Dunnett's multiple comparisons test; $**P < 0.01$, $****P < 0.0001$; ns, not significant.

(F) Representative colony outgrowth from chronological survival assays. Cultures were sampled at the indicated time points, serially diluted ten-fold, and spotted on YPD agar plates to assess viability.

(G, H) Network visualization of enriched Gene Ontology (GO) biological processes among genes downregulated (G) or upregulated (H) following 4-MBA treatment. Nodes represent GO terms, edges indicate functional similarity, and colors denote distinct functional clusters.

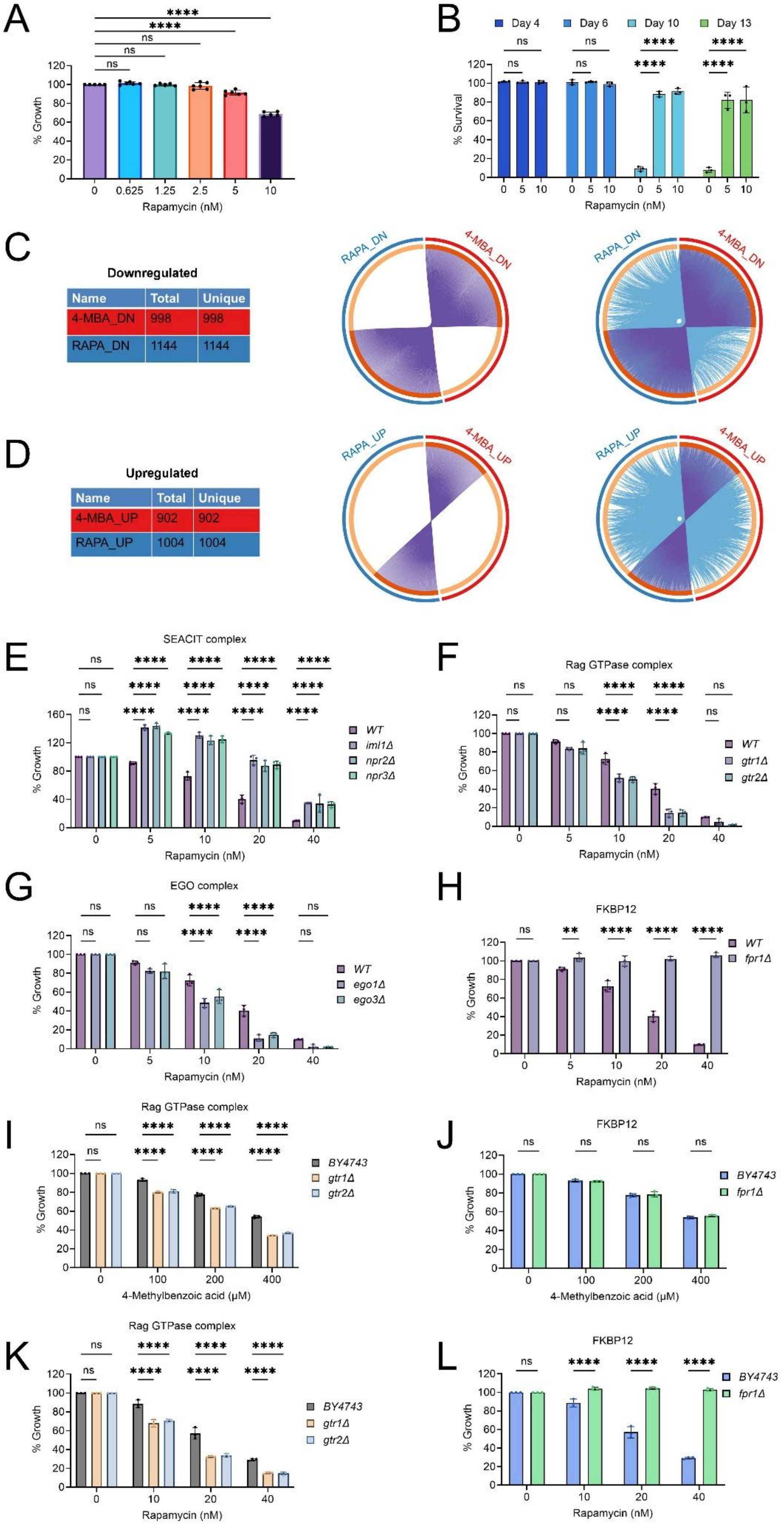
A
% Growth
Rapamycin (nM)
B
Day 4
Day 6
Day 10
Day 13
% Survival
Rapamycin (nM)
C
Downregulated
Name Total Unique
4-MBA_DN 998 998
RAPA_DN 1144 1144
RAPA_DN
4-MBA_DN
D
Upregulated
Name Total Unique
4-MBA_UP 902 902
RAPA_UP 1004 1004
RAPA_UP
4-MBA_UP
E
SEACIT complex
WT
iml1Δ
npr2Δ
npr3Δ
% Growth
Rapamycin (nM)
F
Rag GTPase complex
WT
gtr1Δ
gtr2Δ
% Growth
Rapamycin (nM)
G
EGO complex
WT
ego1Δ
ego3Δ
% Growth
Rapamycin (nM)
H
FKBP12
WT
fpr1Δ
% Growth
Rapamycin (nM)
I
Rag GTPase complex
BY4743
gtr1Δ
gtr2Δ
% Growth
4-Methylbenzoic acid (μM)
J
FKBP12
BY4743
fpr1Δ
% Growth
4-Methylbenzoic acid (μM)
K
Rag GTPase complex
BY4743
gtr1Δ
gtr2Δ
% Growth
Rapamycin (nM)
L
FKBP12
BY4743
fpr1Δ
% Growth
Rapamycin (nM)

**Figure S2. Genetic and transcriptomic comparisons of rapamycin and 4-Methylbenzoic responses, related to Figure 2**

(A) Growth analysis of prototrophic *Saccharomyces cerevisiae* CEN.PK113-7D cells treated with increasing concentrations of rapamycin for 16 h. Data represent mean ± SD (n = 6). Statistical significance was assessed by one-way ANOVA with Dunnett's multiple comparisons test; ****P < 0.0001; ns, not significant.

(B) Chronological survival of CEN.PK113-7D cells treated with increasing concentrations of rapamycin. Viability at each time point is expressed relative to day 3, which was set to 100% as the first measurement following entry into stationary phase. Data represent mean ± SD (n = 3). Statistical significance was assessed by two-way ANOVA with Dunnett's multiple comparisons test; ****P < 0.0001; ns, not significant.

(C, D) Summary tables and Metascape Circos plots showing overlap between gene sets regulated by 4-MBA and rapamycin. Panels display significantly downregulated (C) and upregulated (D) genes, with links indicating shared genes and segment size proportional to gene-set size. Functional overlap highlights shared and condition-specific enriched pathways.

(E–H) Growth assays of CEN.PK113-7D wild-type and TORC1 regulatory mutants treated with increasing concentrations of rapamycin for 16 h, including SEACIT complex components (E), Rag GTPases (F), EGO complex components (G), and FKBP12/Fpr1 (H). Data represent mean ± SD (n = 3). Statistical significance was assessed by two-way ANOVA with Dunnett's multiple comparisons test; **P < 0.01, ****P < 0.0001; ns, not significant.

(I, J) Growth assays of auxotrophic *Saccharomyces cerevisiae* BY4743 wild-type and deletion mutants treated with increasing concentrations of 4-MBA for 16 h, including Rag GTPase mutants (*gtr1Δ*, *gtr2Δ*) (I) and FKBP12/Fpr1 (J). Data represent mean ± SD (n = 3). Statistical significance was assessed by two-way ANOVA with Dunnett's multiple comparisons test; ****P < 0.0001; ns, not significant.

(K, L) Growth assays of BY4743 wild-type and deletion mutants treated with increasing concentrations of rapamycin for 16 h, including Rag GTPase mutants (K) and FKBP12/Fpr1 (L). Data represent mean ± SD (n = 3). Statistical significance was assessed by two-way ANOVA with Dunnett's multiple comparisons test; ****P < 0.0001; ns, not significant.

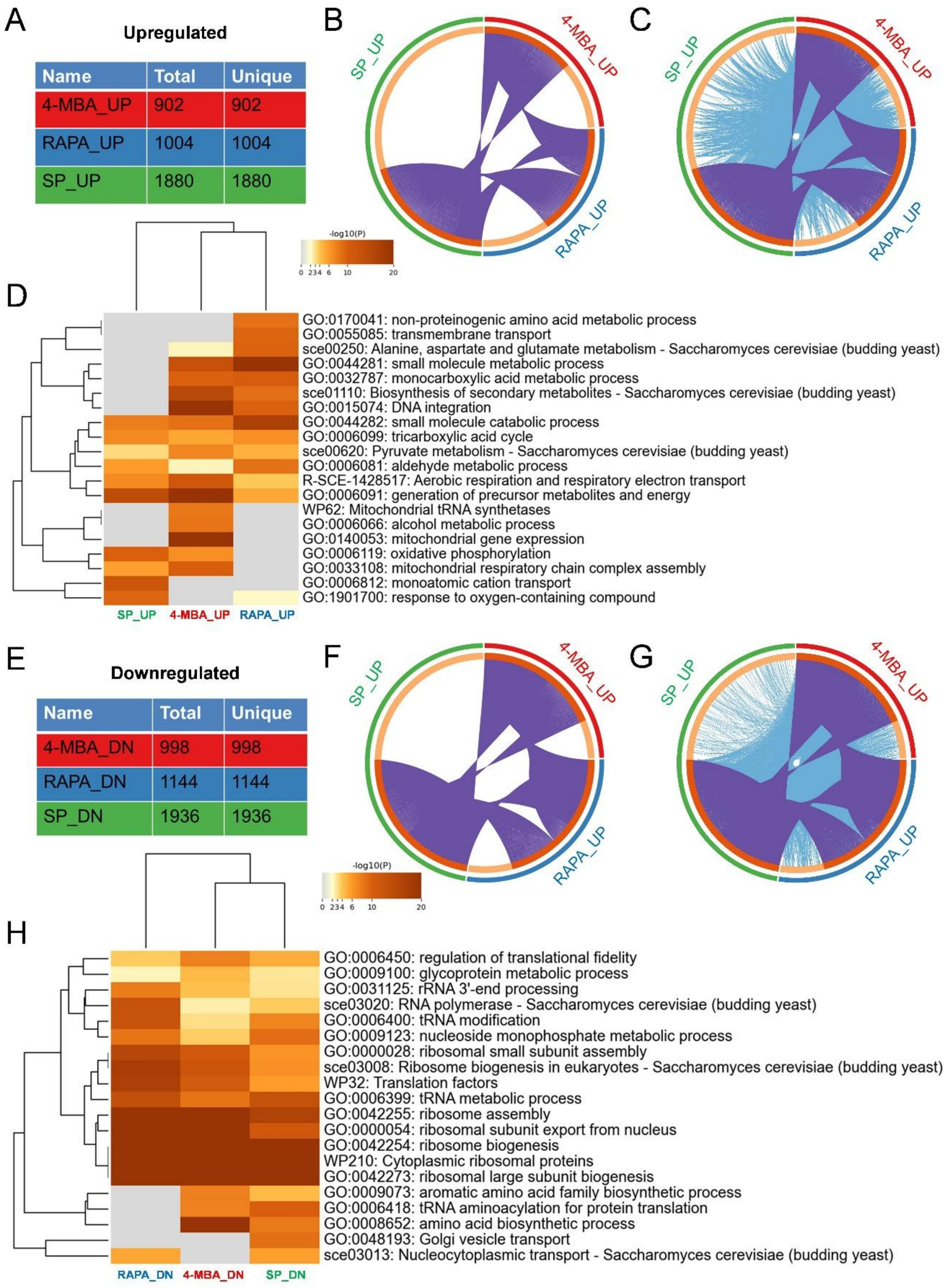


**Figure S3. 4-methylbenzoic acid induces transcriptional programs that converge with rapamycin and stationary-phase adaptive states, related to Figure 3**

(A, E) Summary tables showing the number of genes significantly upregulated (A) or downregulated (E) following treatment with 4-methylbenzoic acid (4-MBA), rapamycin

(RAPA), or during stationary phase (SP), illustrating the extent of transcriptional remodeling across conditions.

(B, F) Metascape Circos plots depicting gene-level overlap among upregulated (B) and downregulated (F) gene sets across the three conditions. Links indicate shared genes, with segment size proportional to gene-set size.

(C, G) Circos plots illustrating overlap among enriched biological processes for upregulated (C) and downregulated (G) gene clusters, highlighting shared functional programs and condition-specific pathways.

(D, H) Heatmaps showing Gene Ontology and pathway enrichment scores comparing functional modules across 4-MBA, rapamycin, and stationary-phase transcriptomes for upregulated (D) and downregulated (H) genes. Color intensity represents $-\log10$(P value), with hierarchical clustering revealing convergence of transcriptional programs associated with adaptive metabolic states.

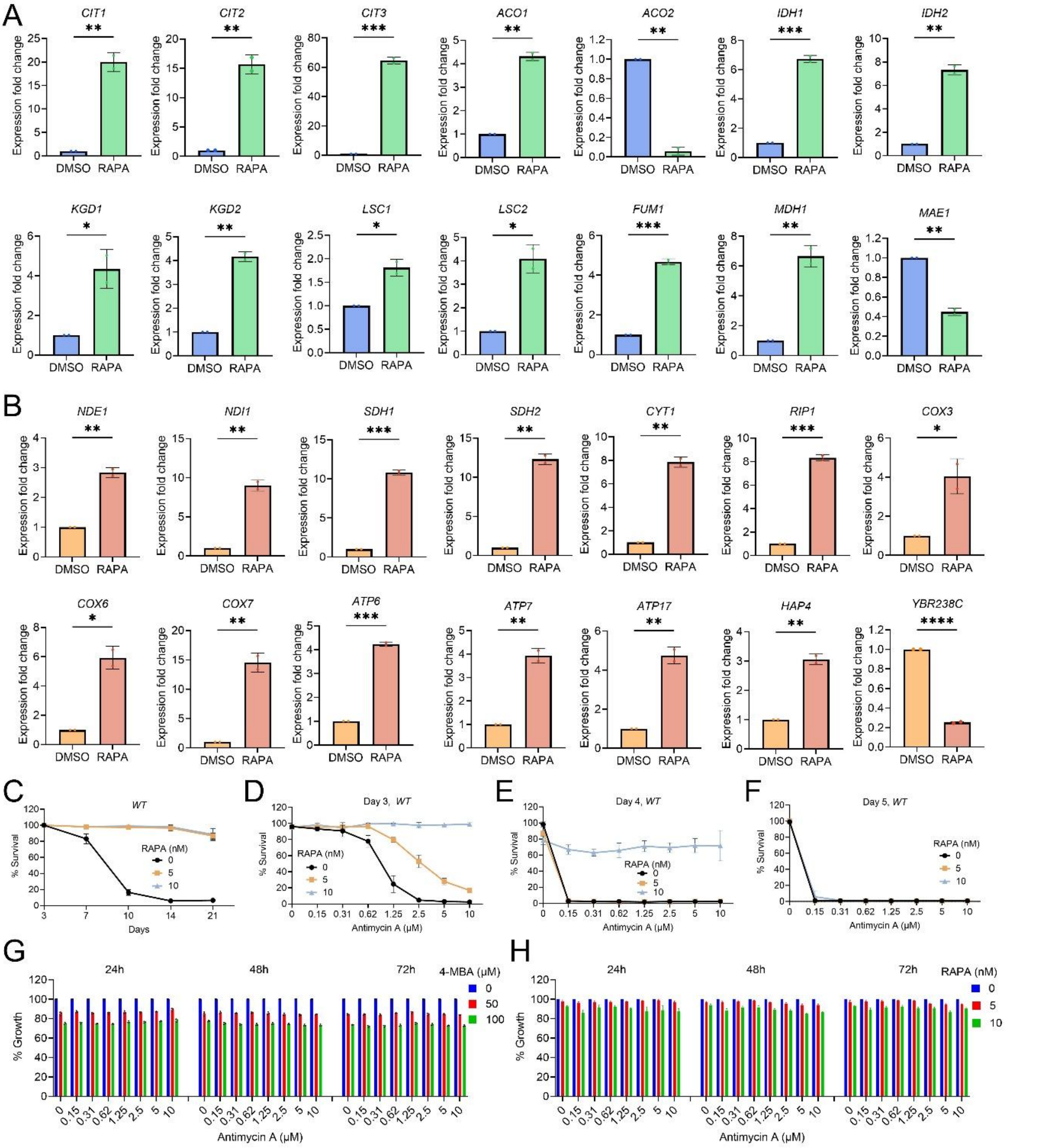


**Figure S4. Rapamycin reinforces mitochondrial programs and reveals respiratory dependence of adaptive survival, related to Figure 3**

(A, B) Quantitative RT–PCR analysis of mitochondrial gene expression in prototrophic *Saccharomyces cerevisiae* CEN.PK113-7D wild-type cells treated with DMSO and rapamycin (200 nM) for 1 h. Panel (A) shows genes involved in the tricarboxylic acid (TCA) cycle, and panel (B) shows genes encoding components of the electron transport chain, including complex I (*NDE1*, *NDI1*), complex II (*SDH1*, *SDH2*), complex III (*CYT1*, *RIP1*), complex IV (*COX3*, *COX6*, *COX7*), and complex V (*ATP6*, *ATP7*, *ATP17*), together with mitochondrial regulators *HAP4* and *YBR238C*. Expression levels were normalized to DMSO-treated controls. Data represent mean ± SD (n = 2). Statistical significance was assessed using two-sided Student's t-tests; *P < 0.05, **P < 0.01, ***P < 0.001, ****P < 0.0001.

(C–F) Chronological survival analyses examining dependence on respiratory function. Wild-type cells were cultured with or without rapamycin in the presence or absence of the respiratory inhibitor antimycin A. Data represent mean ± SD (n = 3).

(G, H) Growth analysis of wild-type cells treated with antimycin A alone or in combination with 4-methylbenzoic acid (4-MBA) (G) or rapamycin (H), assessed at 24 h, 48 h, and 72 h. Growth is expressed relative to untreated controls. Data represent mean ± SD (n = 3).

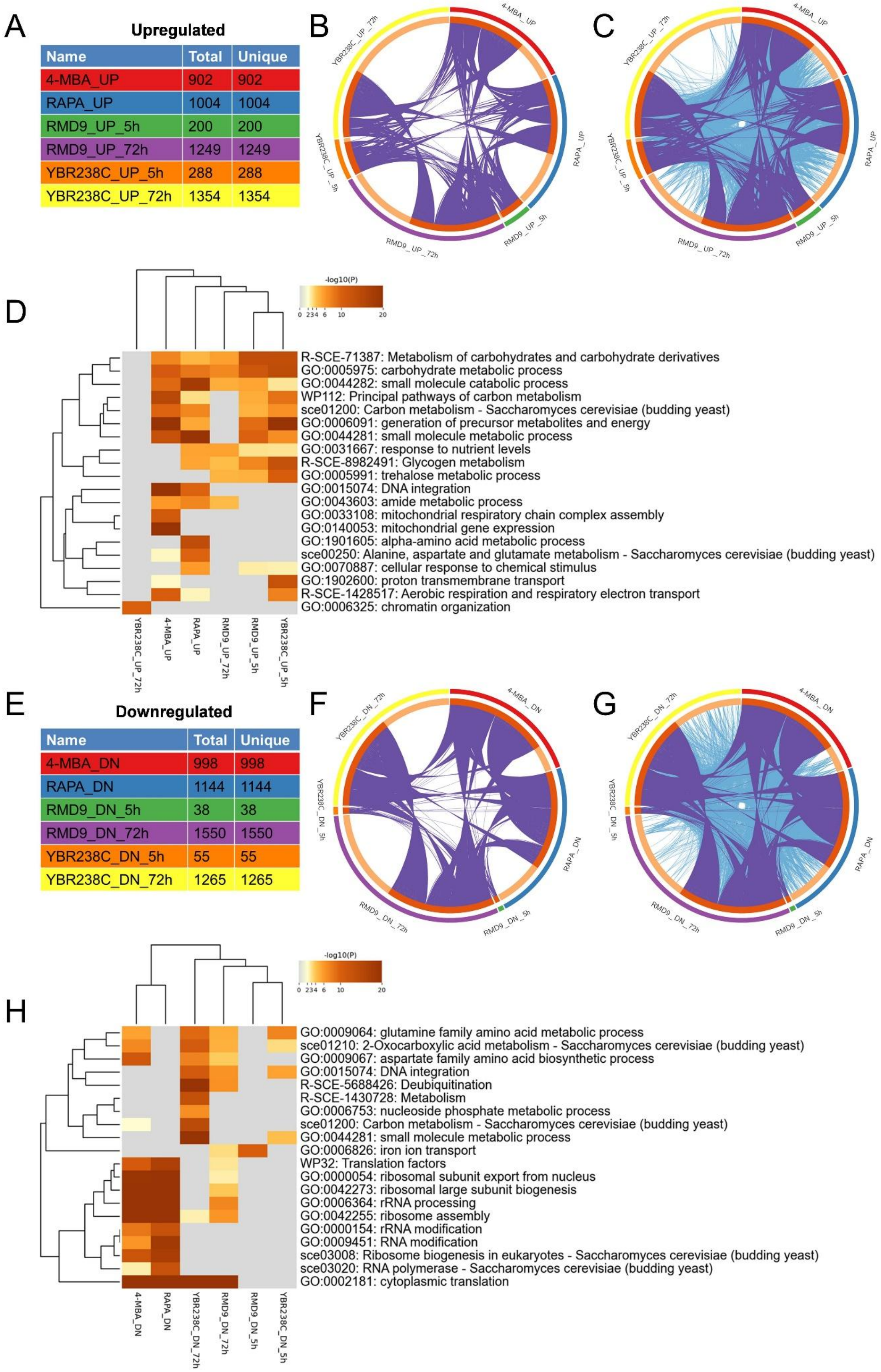

**Upregulated**

| Name | Total | Unique |
|---|---|---|
| 4-MBA_UP | 902 | 902 |
| RAPA_UP | 1004 | 1004 |
| RMD9_UP_5h | 200 | 200 |
| RMD9_UP_72h | 1249 | 1249 |
| YBR238C_UP_5h | 288 | 288 |
| YBR238C_UP_72h | 1354 | 1354 |

**Downregulated**

| Name | Total | Unique |
|---|---|---|
| 4-MBA_DN | 998 | 998 |
| RAPA_DN | 1144 | 1144 |
| RMD9_DN_5h | 38 | 38 |
| RMD9_DN_72h | 1550 | 1550 |
| YBR238C_DN_5h | 55 | 55 |
| YBR238C_DN_72h | 1265 | 1265 |

**Figure S5. Transcriptomic comparisons across chemical perturbations and mitochondrial regulatory states, related to Figure 4**

(A, E) Summary tables showing the number of genes significantly upregulated (A) or downregulated (E) following treatment with 4-methylbenzoic acid (4-MBA) or rapamycin (RAPA), and in mitochondrial regulatory mutants (*rmd9Δ* and *ybr238cΔ*) at early (5 h) and late (72 h) time points.

(B, F) Metascape Circos plots depicting gene-level overlap among upregulated (B) and downregulated (F) gene sets across conditions. Links indicate shared genes, and segment size reflects gene-set size.

(C, G) Circos plots illustrating overlap among enriched biological processes for upregulated (C) and downregulated (G) gene clusters, highlighting shared functional programs and condition-specific pathways.

(D, H) Heatmaps showing Gene Ontology and pathway enrichment scores comparing functional modules across transcriptomes for upregulated (D) and downregulated (H) genes. Color intensity represents −log10(P value), with hierarchical clustering revealing convergence of transcriptional programs associated with adaptive metabolic configurations.

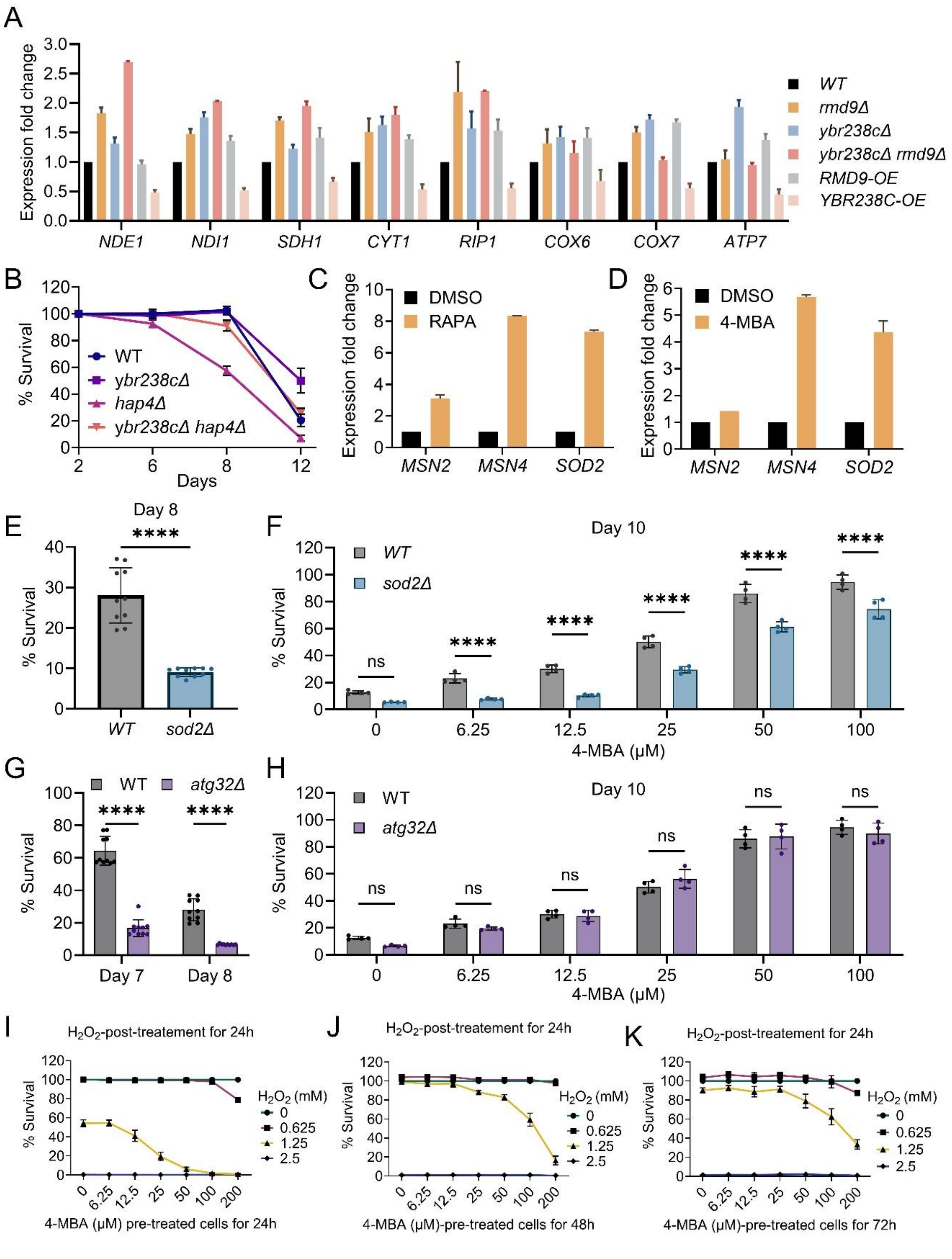


**Figure S6. Adaptive stress responses and trade-offs associated with mitochondrial state, related to Figure 4**

(A) Quantitative RT–PCR analysis of mitochondrial gene expression across indicated genotypes during exponential growth. Expression levels were normalized to wild-type controls. Data represent mean ± SD (n = 2).

(B) Chronological survival analysis of prototrophic *Saccharomyces cerevisiae* CEN.PK113-7D wild-type and mitochondrial regulatory mutants (*ybr238cΔ*, *hap4Δ*, and double mutant) cultured in synthetic defined medium. Viability is expressed relative to the initial stationary phase measurement. Data represent mean ± SD (n = 6).

(C, D) Quantitative RT–PCR analysis of stress response regulators (*MSN2*, *MSN4*, *SOD2*) in exponential-phase wild-type cells treated with rapamycin (200 nM) (C) or 4-methylbenzoic acid (4-MBA; 200 μM) (D) for 1 h. Expression levels were normalized to DMSO-treated controls. Data represent mean ± SD (n = 2).

(E–H) Chronological survival analysis of wild-type, *sod2Δ*, and *atg32Δ* cells cultured without treatment (E, G) or in the presence of 4-methylbenzoic acid (4-MBA) (F, H). Viability is expressed relative to the initial stationary phase measurement. Data represent mean ± SD (untreated n = 10; 4-MBA n = 6). Statistical significance was assessed using an unpaired two-tailed Student's t test for untreated conditions and two-way ANOVA with Dunnett's multiple comparisons test for treatment comparisons; ****P < 0.0001; ns, not significant.

(I–K) Chronological survival of wild-type cells preincubated with 4-MBA for 24 h (I), 48 h (J), or 72 h (K) in synthetic defined medium followed by exposure to $H_2O_2$.

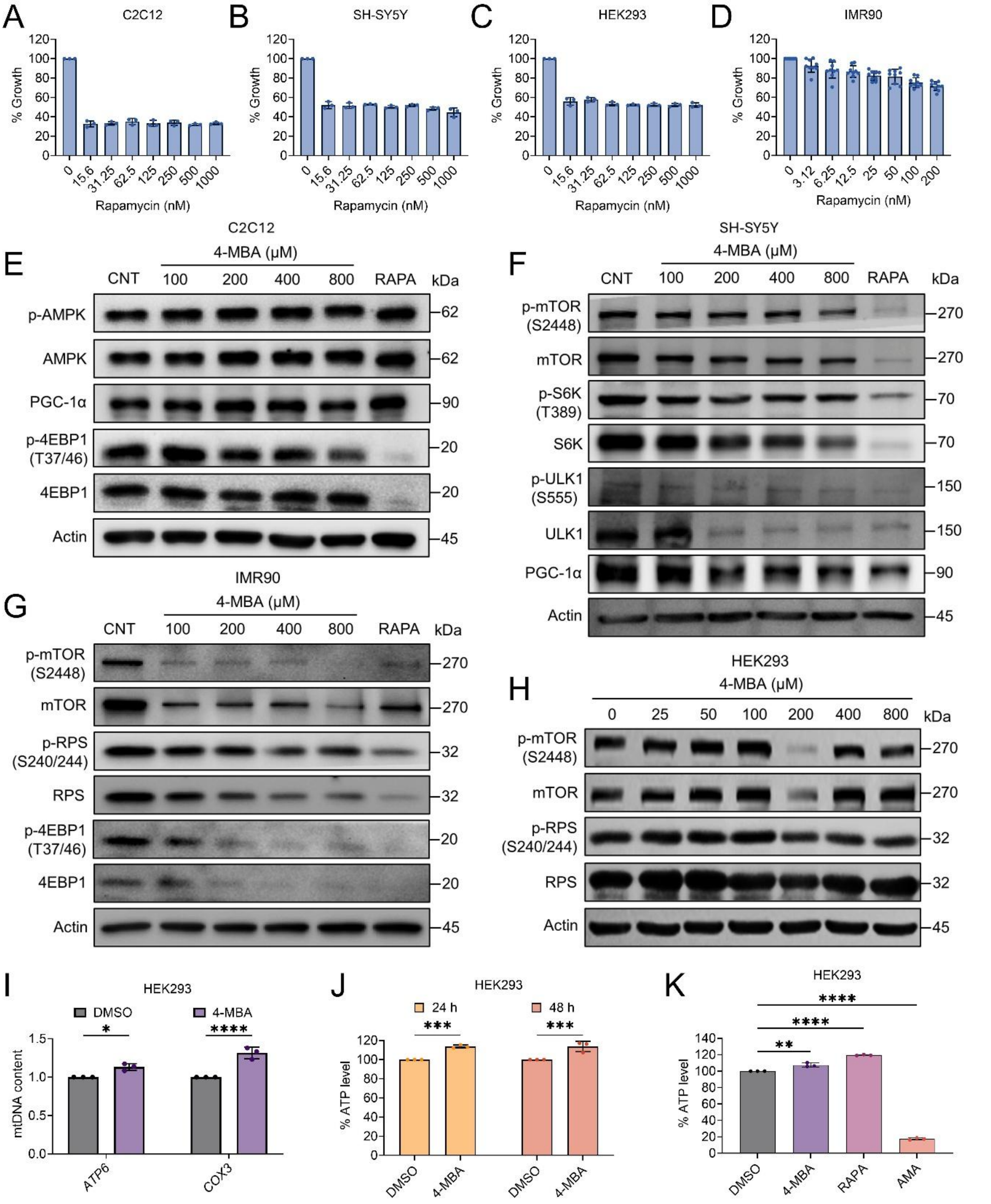


**Figure S7. Mammalian validation of growth, signaling, and mitochondrial responses to TORC1 modulation, related to Figures 5 and 6**

(A–D) Growth analysis of mammalian cells treated with rapamycin for 48 h, including (A) mouse myoblasts (C2C12), (B) human neuroblastoma cells (SH-SY5Y), (C) human embryonic kidney cells (HEK293), and (D) human lung fibroblasts (IMR90). Cell growth was quantified using the CCK-8 assay and expressed as a percentage relative to DMSO-treated controls. Data represent mean ± SD (n ≥ 3).

(E–H) Representative immunoblots showing signaling responses in (E) C2C12, (F) SH-SY5Y, (G) IMR90, and (H) HEK293 cells treated with DMSO, 4-methylbenzoic acid (4-MBA), or rapamycin (100 nM) for 1 h. Blots show phosphorylated and total forms of mTOR, S6K, ribosomal protein S6 (RPS6), and 4EBP1, along with phosphorylated and total AMPK and ULK1, and PGC-1α as indicated. β-Actin was used as a loading control.

(I) Relative mitochondrial DNA (mtDNA) content in HEK293 cells treated with 4-MBA (50 μM). mtDNA levels were quantified by qPCR of mitochondrial genes (*ATP6* and *COX3*) and normalized to the nuclear gene *GAPDH*. Data represent mean ± SD (n = 4). Statistical significance was assessed by two-way ANOVA with Dunnett's multiple comparisons test; **P < 0.01.

(J) Intracellular ATP levels in HEK293 cells treated with 4-MBA (200 μM) for 24 h and 48 h. Data represent mean ± SD (n = 3). Statistical significance was determined by two-way ANOVA with Šídák's multiple comparisons test; ***P < 0.001.

(K) ATP levels in HEK293 cells treated with 4-MBA (100 μM), rapamycin (100 nM), or antimycin A (10 μM) for 6 h. Data represent mean ± SD (n = 3). Statistical significance was assessed by one-way ANOVA with Dunnett's multiple comparisons test; **P < 0.01, ****P < 0.0001.

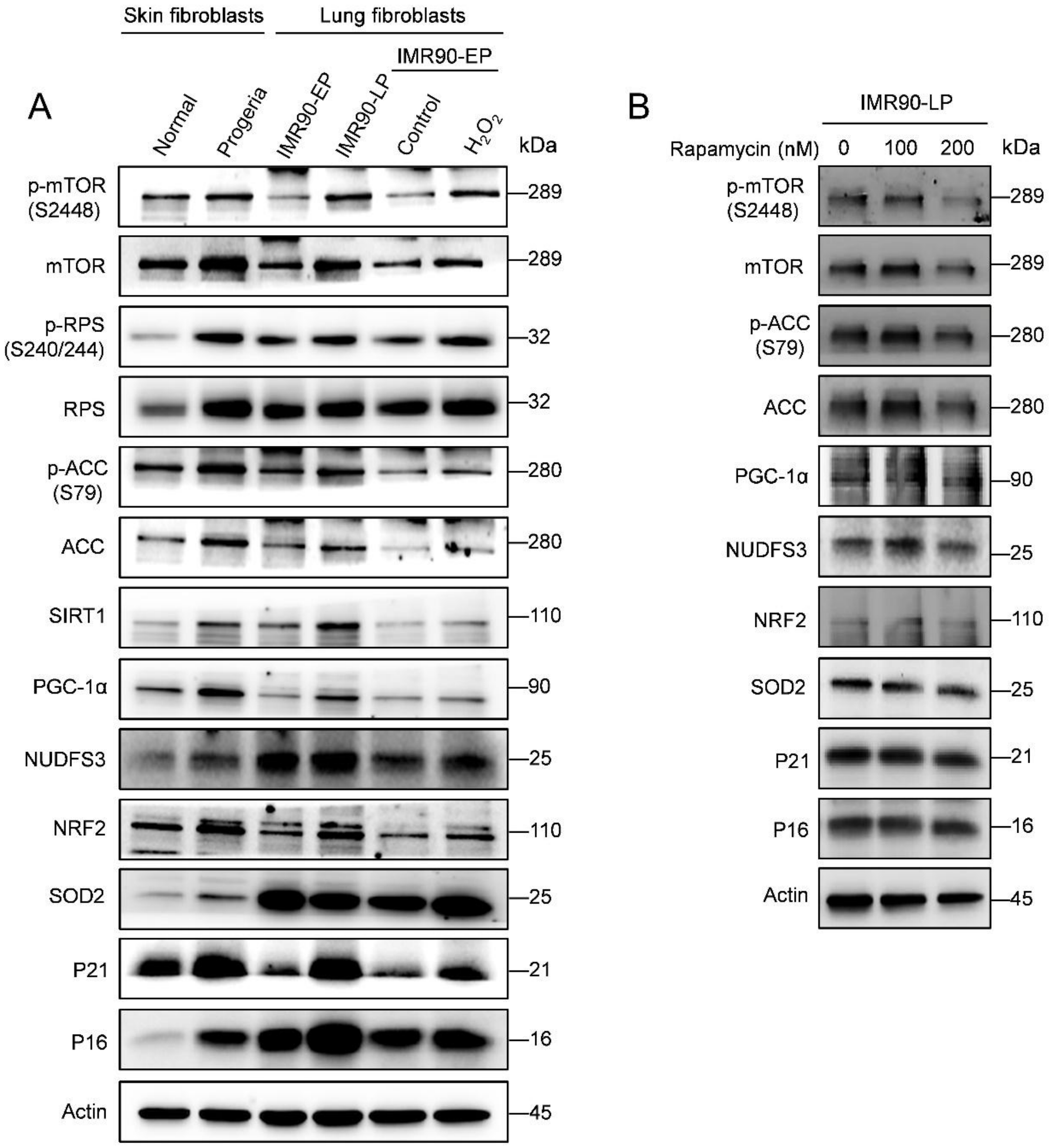


**Figure S8. Validation of signaling states and rapamycin responses across physiological contexts, related to Figure 7**

(A) Replicate immunoblot analysis comparing signaling states across physiological contexts. Human skin fibroblasts from normal donors (AG03512) and Hutchinson–Gilford progeria syndrome (HGPS; AG03513), both at passage 14, were analyzed alongside IMR90 fibroblasts at early passage (EP-15), late passage (LP-25), and early-passage IMR90 cells exposed to hydrogen peroxide ($H_2O_2$; 200 μM, 4 h). Blots show phosphorylated and total mTOR (Ser2448), ribosomal protein S6 (Ser240/244), acetyl-CoA carboxylase (Ser79), and markers of mitochondrial regulation (SIRT1, PGC-1α, NDUFS3), antioxidant response (NRF2, SOD2), and cell cycle checkpoints (p21, p16). β-Actin was used as a loading control.

(B) Immunoblot analysis of late-passage IMR90 fibroblasts (LP-26) treated with rapamycin for 2 h. Blots show phosphorylated and total mTOR (Ser2448), ribosomal protein S6 (Ser240/244), acetyl-CoA carboxylase (Ser79), and markers of mitochondrial regulation (PGC-1α, NDUFS3), antioxidant response (NRF2, SOD2), and checkpoint proteins (p21, p16). β-Actin served as a loading control.

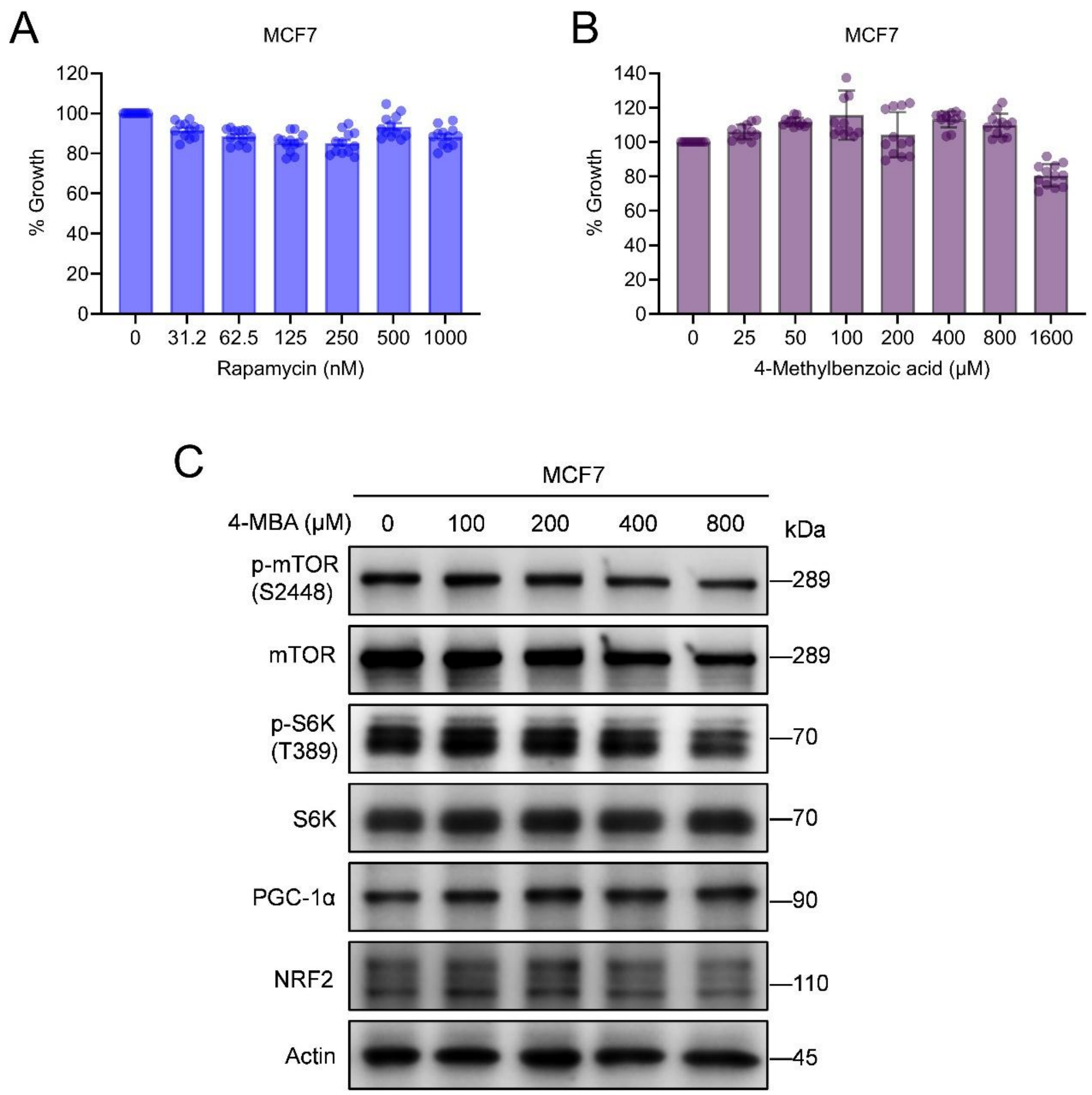


**Figure S9. 4-MBA modulates growth and signaling in MCF7 cells, related to Figure 7.**

(A,B) Growth analysis of MCF7 cells treated with (A) rapamycin or (B) 4-methylbenzoic acid (4-MBA) for 48 h. Cell viability was quantified using the CCK-8 assay and expressed as a percentage relative to DMSO-treated controls. Data represent mean ± SD (n ≥ 3).

(C) Representative immunoblots showing signaling responses following 24 h treatment with the indicated concentrations of 4-MBA. Blots show phosphorylated and total mTOR and S6K, as well as PGC-1α and NRF2. β-Actin served as a loading control.
